# Spatial proteomics reveals prefrontal circuit diversity in socioemotional behaviour

**DOI:** 10.64898/2026.02.08.704617

**Authors:** Yuki Ito, Sayaka Nagamoto, Sawako Uchiyama, Kohei Onishi, Junpei Matsubayashi, Akari Fukuda, Mizuki Funakoshi, Satoko Hattori, Kosei Takeuchi, Mikiko Kudo, Takahiro Masuda, Yuichi Hiraoka, Yuta Kohro, Tomomi Shimogori, Makoto Tsuda, Hidetaka Kosako, Tetsuya Takano

## Abstract

Understanding how molecular diversity across long-range neural circuits governs brain function remains a central challenge in neuroscience. Here, we developed a projection-specific spatial proteomics approach and revealed robust presynaptic molecular divergence across six projection-defined pathways of the medial prefrontal cortex (mPFC), including efferent projections to the basolateral amygdala (BLA), nucleus accumbens (NAc), thalamus (Thal), hypothalamus (HT) and cortex (CTX), as well as the afferent projection from the BLA to the mPFC. Among these pathways, we identify BLTP2 (KIAA0100), a previously uncharacterized transmembrane protein, as highly enriched in the projection from the mPFC to the BLA. BLTP2 localizes to excitatory presynaptic terminals and is enriched at synapses that are activated during memory formation. Loss of BLTP2 impairs synaptic structure and transmission, and reduces activity-dependent remodelling, resulting in selective deficits in contextual fear memory, anxiety-related behavior, and social behavior. Mechanistically, BLTP2 promotes presynaptic assembly by recruiting Neurexin 1. These findings reveal projection-specific presynaptic molecular diversity and provide mechanistic insights into circuit-level vulnerabilities in neuropsychiatric disorders.

## Main

Long-range neural projections form synaptic connections with specific target regions and thereby route information across distributed brain circuits ^1^. Anatomical and physiological studies indicate that these projections are not uniform but instead form synapses with distinct structural and functional properties, which supports selective information processing in defined neural pathways ^2–5^. A central challenge in neuroscience is to define how projection-specific molecular architectures, particularly at presynaptic terminals, contribute to this functional diversity and to behavioural control. Recent transcriptomic and single-cell RNA sequencing studies have revealed extensive neuronal heterogeneity and have shown that neurons in regions such as the medial prefrontal cortex (mPFC) exhibit target-dependent gene expression profiles ^6–12^. However, transcript abundance alone does not reliably predict protein composition at the synapse, because local translation, trafficking and degradation generate compartment-specific proteomes that can diverge from the somatic transcriptome ^13–16^. Direct proteomic profiling of presynaptic terminals in defined projection pathways is therefore essential to establish the molecular architectures that underlie projection-specific synaptic identity and to understand how these architectures relate to circuit function and disease vulnerability ^17,18^.

The mPFC exerts top-down control in the brain through long-range collateral projections to anatomically and functionally distinct targets such as the basolateral amygdala (BLA), nucleus accumbens (NAc), thalamus (Thal), hypothalamus (HT) and cortical regions (CTX) ^4,19–24^. Each of these projection-defined neural pathways contributes to distinct behavioural functions. Moreover, distinct behavioural roles have been identified in the reciprocal connectivity between the mPFC and BLA, with the mPFC-to-BLA (mPFC–BLA) projection playing a central role in regulating fear memory, anxiety-like behaviour and social interaction, whereas the BLA–mPFC projection mediates fear extinction and affective salience ^25–29^. These functionally segregated pathways are also differentially implicated in neuropsychiatric and neurodevelopmental disorders, including autism spectrum disorder (ASD), schizophrenia, post-traumatic stress disorder, major depression and social anxiety ^30,31^. Despite this behavioural specificity and clinical relevance, the presynaptic molecular architecture of the mPFC–BLA pathway remains poorly defined ^32^.

Proteomic analysis of defined projections *in vivo* remains technically challenging. Conventional approaches based on tissue homogenization or synaptosome purification lack spatial specificity and average signals across many cell types and pathways, which prevents isolation of presynaptic proteomes from individual projection pathways in intact circuits ^33,34^. Proximity-dependent biotinylation approaches such as BioID have enabled spatially restricted labelling of proteins in neurons using engineered biotin ligases (e.g., BirA*R118G, TurboID, ultraID) ^35–37^. These enzymes covalently tag neighbouring proteins with biotin, which can then be enriched by streptavidin pulldown and analysed by mass spectrometry. We have previously reported the successful application of BioID strategies for profiling the proteomes of specific cell types and cell–cell interfaces in the intact brain ^33,34,38,39^. However, existing approaches still lack a broadly applicable *in vivo* solution that simultaneously provides pathway-level spatial precision, deep proteomic coverage, and compatibility with routine workflows.

To address this gap, we asked whether presynaptic terminals in projection defined pathways of the mPFC possess distinct protein architectures that could account for pathway specific synaptic properties, behavioural specialisation and disease relevance. We therefore mapped pathway resolved presynaptic proteomes across six major mPFC centred circuits, including outputs to the BLA, NAc, Thal, HT and CTX, as well as the reciprocal input from the BLA to the mPFC. This analysis reveals marked molecular divergence across projection defined synapses and identifies pathway enriched programmes linked to vesicle trafficking, cytoskeletal organisation, synaptic adhesion and intracellular signalling. Focusing on the mPFC to BLA pathway, we uncover BLTP2 (KIAA0100) as a selectively enriched presynaptic transmembrane protein and show that it promotes presynaptic assembly through recruitment of Neurexin 1, thereby supporting synaptic structure, excitatory transmission and activity dependent remodelling. Together, these findings define projection specific presynaptic molecular identity as an organising principle of long range mPFC circuits and provide a mechanistic entry point into circuit selective vulnerability in neuropsychiatric disorders.

## Results

### Development of ProX-ID enables projection-specific presynaptic proteome profiling in the reciprocal mPFC–BLA neural pathways

Building on our previous *in vivo* applications of BioID ^19,20,24,25^, we developed Projection-eXclusive BioID (ProX-ID), a proximity-labelling system designed for neural pathway-specific proteomic profiling of presynaptic terminals in the bidirectional mPFC**–**BLA projections (Fig. 1a–b). ProX-ID consists of the ultraID enzyme fused to the palmitoylation sequence of growth-associated protein 43 (GAP43), enabling its localization to presynaptic membranes (Fig. 1a). We first validated ProX-ID activity *in vitro* using cultured neurons at 14 days *in vitro* (DIV14), which showed robust biotinylation signals upon biotin treatment (Extended Data Fig.1a–b). To assess *in vivo* labelling, we transduced an adeno-associated virus (AAV-CaMKII-ProX-ID) into either the prelimbic (PrL) region of the mPFC or the BLA of six-week-old mice (Fig. 1b).

**Figure 1.**
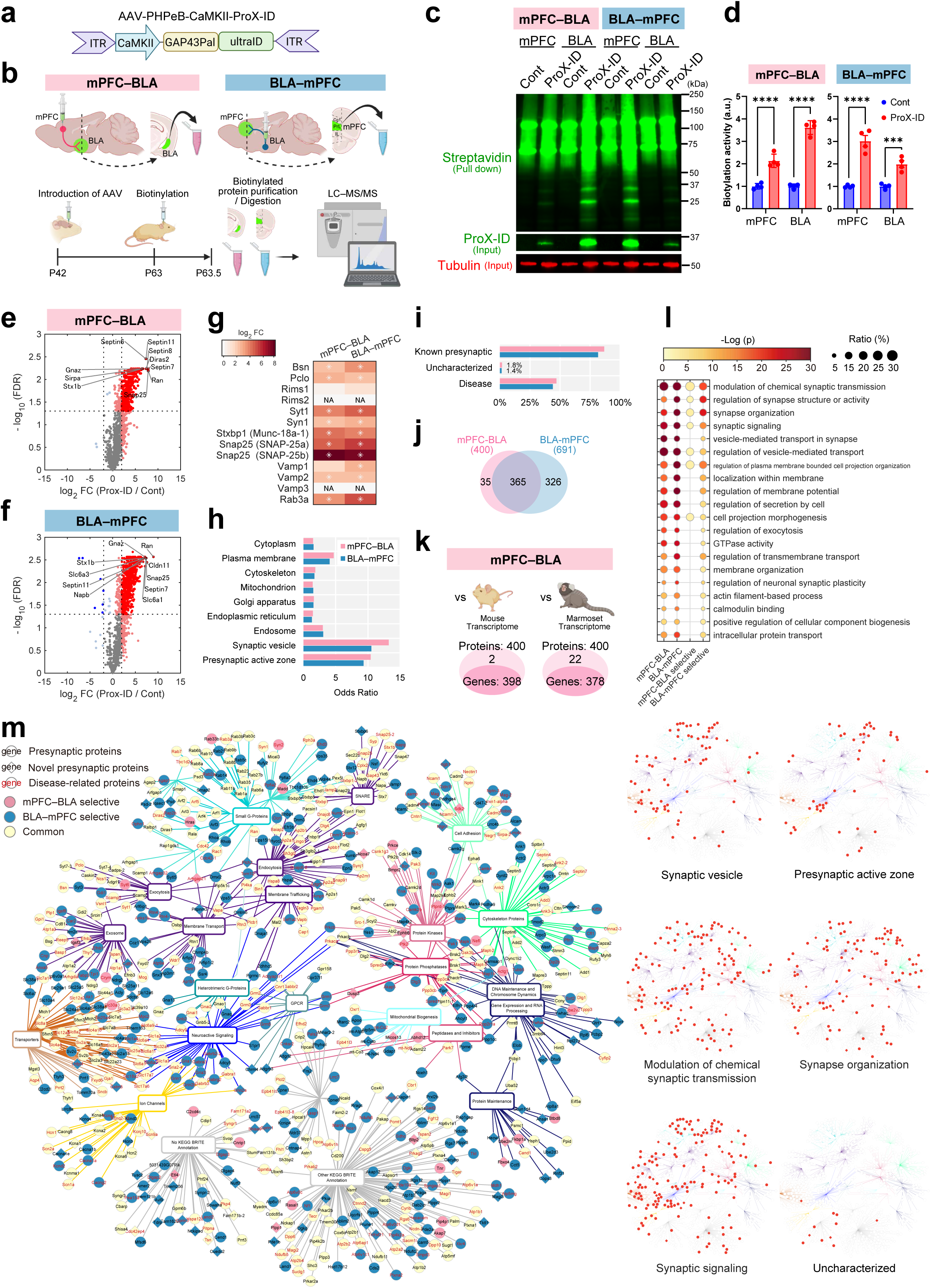
Establishment and validation of ProX-ID for projection-specific presynaptic proteome profiling in mPFC–BLA neural pathways. **a,** Schematic of the ProX-ID method. The palmitoylation sequence of GAP43 is fused to ultraID to enable proximity-dependent biotinylation of presynaptic proteins in mPFC–BLA and BLA–mPFC projections. **b,** Experimental design of *in vivo* ProX-ID labelling. **c,** Immunoblot analysis of biotinylated proteins in the mPFC–BLA and BLA–mPFC neural pathways. **d,** Quantification of streptavidin signals in the mPFC–BLA and BLA–mPFC neural pathways. Data are presented as mean ± s.e.m.; *n* = 4 mice per experimental group. Two-way ANOVA followed by Bonferroni *post hoc* test. \*\*\*\**P* < 0.0001; \*\*\**P* < 0.001. Results for the selected pairwise comparisons are shown. **e**–**f,** Volcano plots for mPFC–BLA (**e**) and BLA–mPFC (**f**) neural pathways. Dashed lines indicate FDR = 0.05 (horizontal) and log₂ fold change = −2 and 2 (vertical). Red dots represent significantly enriched proteins (FDR < 0.05, log₂ fold change > 2). Gene symbols indicate the top 10 proteins with the largest fold changes. **g,** Heatmap of normalized log₂ biotinylation levels (ProX-ID vs. control) for representative presynaptic proteins. Asterisks indicate proteins significantly enriched in ProX-ID samples (FDR < 0.05, log_2_ fold change > 2, ProX-ID vs. control). NA indicates missing values. **h,** Odds ratios for selected Gene Ontology (GO) Cellular Component terms (see Materials and Methods). **i,** Proportion of known presynaptic, neurological disease-associated and functionally uncharacterized proteins among the neural pathway-enriched proteins. **j,** Venn diagram showing the overlap of significantly enriched proteins between the mPFC–BLA and BLA–mPFC neural pathways. Numbers indicate protein numbers. **k,** Overlap between mPFC–BLA-enriched proteins and transcriptomes from mouse (left) and marmoset (right). Numbers indicate protein numbers. **l,** Top 20 enriched GO terms for Biological Process and Molecular Function. mPFC–BLA selective and BLA–mPFC selective describe proteins selectively enriched in the mPFC–BLA and BLA–mPFC neural pathway, respectively. **m,** Protein–term network based on KEGG BRITE annotations (left). Nodes represent proteins and functional terms, with edges indicating associations based on BRITE functional categories. Proteins annotated for the selected category are highlighted (right).

Following a three-week expression period, mice were administered a single subcutaneous injection of biotin (24 mg/kg), and brains were collected 12 h later for analysis (Fig. 1b). Immunohistochemical and immunoblot analysis revealed that biotinylated proteins were prominent in the BLA when ProX-ID was expressed in the PrL region of the mPFC (Fig. 1c and d, Extended Data Fig.1c and d). Conversely, when ProX-ID was expressed in the BLA, biotinylated proteins were observed in the PrL region of the mPFC (Fig. 1c and d, Extended Data Fig.1e and f). Super-resolution stimulated emission depletion (STED) microscopy revealed that biotinylated proteins were selectively localized at mCherry-labelled axonal terminals in both neural pathways, confirming the specificity of the labelling (Extended Data Fig. 1g). Moreover, the biotinylated proteins were highly enriched in excitatory synapses, as evidenced by their colocalization with Homer1 and VGLUT1 (Extended Data Fig. 1h).

To further characterize the molecular composition of these projection-specific presynaptic terminals, we conducted quantitative spatial proteomic profiling of synapses in the defined mPFC–BLA and BLA–mPFC neural pathways using streptavidin pull-down followed by mass spectrometry. We detected 1,402 unique proteins from mPFC–BLA and BLA–mPFC projection neurons *in vivo* (Extended Data Table 1). In three independent experiments, 400 proteins were significantly enriched in the mPFC–BLA neural pathway, while 691 proteins were significantly enriched in the BLA–mPFC neural pathway (log_2_ fold change (FC) > 2.0, FDR < 0.05; Fig. 1e and f, Extended Data Table 1 and 2). Notably, several known presynaptic proteins including Snap25, Septin7/11, Bsn, Syt1, Syn1, Munc18 and Vamp were robustly recovered in both pathways, supporting the sensitivity and specificity of ProX-ID *in vivo* (Fig. 1e, f and g) ^40–42^. Gene Ontology (GO) Cellular Component annotations showed a predominant association of identified proteins to synaptic vesicles (GO:0008021) and the presynaptic active zone (GO:0048786) compared to other cellular compartments (Fig. 1h, Extended Data Fig. 2a and b). Comparison with the SynaptomeDB database indicated more than 80% overlap with a previously reported presynaptic proteome ^43^ (Fig. 1i, Extended Data Table 2). A further 1–2% of proteins were functionally uncharacterized (see Materials and Methods), and 43–47% were associated with neurological or psychiatric disorders (Fig. 1i, Extended Data Tables 2 and 3), supporting the existence of previously unrecognized synaptic components and implicating projection-enriched proteins in disease.

Neural pathway comparisons revealed distinct signatures, with 35 proteins selectively enriched in the mPFC–BLA projection, 326 in the BLA–mPFC projection, and 365 shared (Fig. 1j, Extended Data Table 2). Spatial transcriptomic datasets supported anatomical fidelity and cross-species conservation. Of the 400 proteins enriched in the mPFC–BLA projection, 398 (99.5%) corresponded to gene expression in layer 2/3 of mouse mPFC (Fig. 1k; Extended Data Fig. 2c, 3a), and 378 (95%) had orthologues expressed in marmoset mPFC (Fig. 1k; Extended Data Fig. 2d, 2e, 3b). GO enrichment analysis with Metascape ^44^ highlighted shared core programmes required for synaptic communication across directions, including synaptic transmission (GO:0050804), synapse organization (GO:0050808), synaptic signalling (GO:0099536), vesicle-mediated transport in synapse (GO:0099003) and regulation of membrane potential (GO:0042391) (Fig. 1l, Extended Data Table 2). Protein–term networks based on the KEGG BRITE database indicated that the mPFC–BLA neural pathway-selective proteome comprises synaptic vesicle proteins including exocytosis and endocytosis (e.g., Syt6, Ncoa7), synaptic organization proteins including cytoskeletal proteins (Actg1, and Nefl), cell adhesion proteins (Sdk2 and Ptprd), synaptic transmission related proteins including transporters (Slc30a3), small GTPases and associated proteins (e.g., Rab33b, Syn2), and synaptic signalling molecules (e.g., Prkce, Ephb6, Bmpr2) (Fig. 1m, Supplemental Fig. 1). Gene expression mapping showed that proteins for exocytosis, the SNARE complex, cell adhesion and cytoskeletal organization were relatively highly expressed in BLA-projecting mPFC neurons, in partial concordance with our proteomics (e.g., Snap25) (Extended Data Fig. 3a). Interestingly, homologues of many of the most highly expressed proteins in the marmoset mPFC are associated with human neurological and psychiatric disorders, suggesting conservation of disease-relevant molecular features across species (Extended Data Fig. 3b). Together, these findings demonstrate that ProX-ID enables neural pathway-specific presynaptic proteomic profiling and reveals distinct molecular architectures between the directionally distinct projections of the mPFC–BLA and BLA–mPFC pathways.

### Projection-specific proteomic profiling reveals molecular heterogeneity across mPFC output neural pathways

Building on the projection-specific proteomic profiles established for the mPFC–BLA and BLA–mPFC neural pathways (Fig. 1), we next extended our analysis to investigate projection-specific synaptic proteomes across multiple mPFC output neural pathways (Fig. 2a). In the PrL-targeted same brains, immunohistochemistry and immunoblotting revealed robust biotinylation in the nucleus accumbens (NAc), mediodorsal thalamus (MD; hereafter referred to as Thal), lateral hypothalamic area (LHA; hereafter referred to as HT) and ipsilateral motor and insular cortices (collectively referred to as CTX) (Fig. 2a–c, Extended Data Fig. 4a and b). To comprehensively characterize the synaptic proteomes of these mPFC output neural pathways, we performed ProX-ID-based quantitative spatial proteomic profiling *in vivo* (Fig. 2a). Across four independent experiments, we detected 3,125 biotinylated peptides corresponding to 1,324 unique biotinylated proteins from mPFC–NAc, mPFC–Thal, mPFC–HT and mPFC–CTX projection neurons. Among these, 815, 491, 573 and 176 proteins were significantly enriched in each neural pathway (log_2_ fold change (FC) > 2.0, FDR < 0.05; Fig. 2d–g, Extended Data Table 1). Enriched proteomes included presynaptic molecules (Ap2a1, Atp6v1a), cytoskeletal proteins (Tbcb, Myo18a) and small GTPase-related proteins (Diras2, Rasl2-9). We also detected the carbonyl reductase Cbr1, previously implicated in spatial memory formation ^45^. Consistently, numerous canonical presynaptic proteins were confirmed in our dataset (Fig. 2h, Extended Data Fig. 4c), and several showed neural pathway-dependent abundance and isoform usage including SNAP-25a, SNAP-25b, Vamp2 and Vamp3 (Fig. 2h).

**Figure 2.**
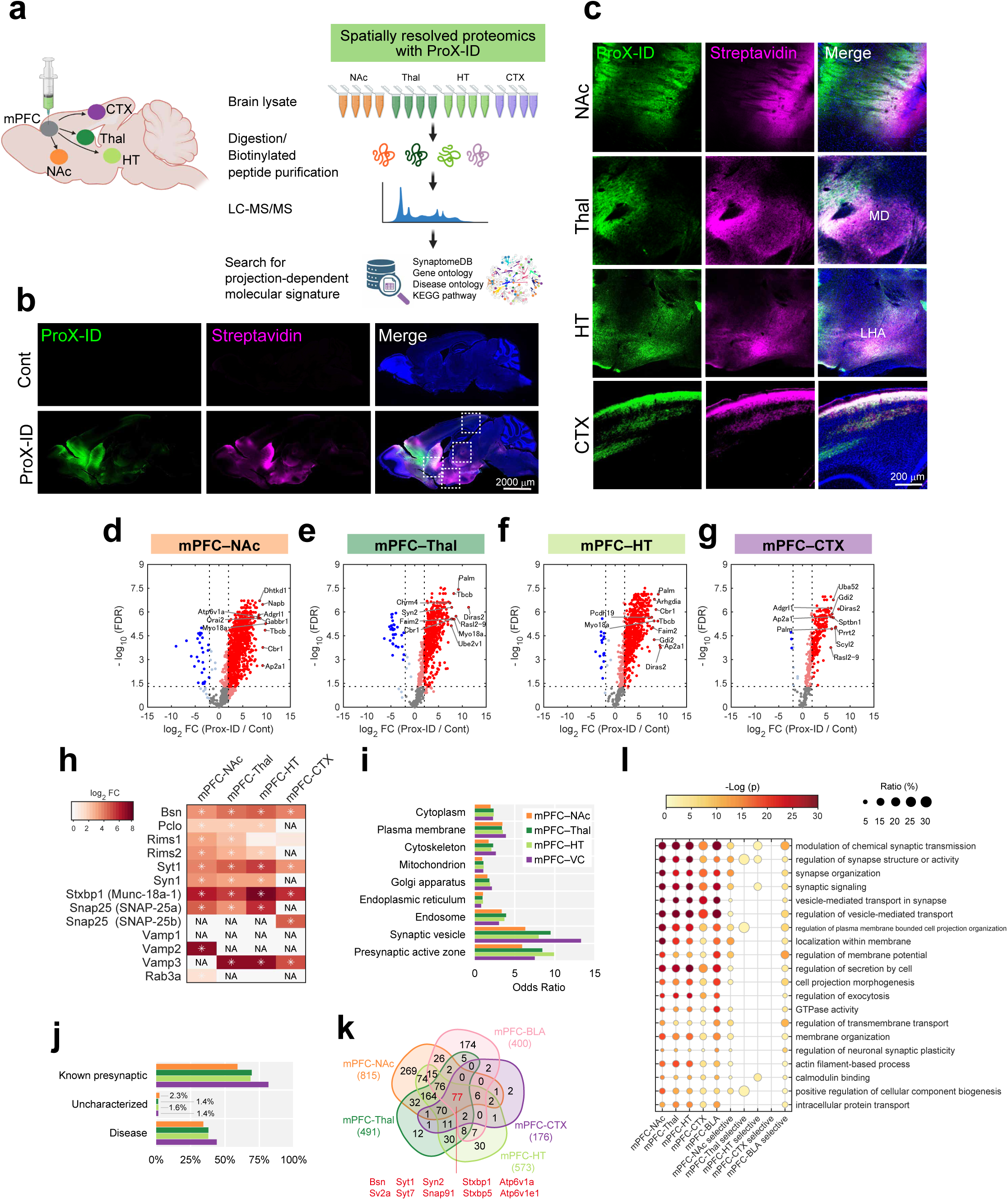
Molecular profiling of mPFC outputs reveals projection-specific presynaptic landscapes. **a,** Schematic of the ProX-ID method targeting multiple mPFC projection targets. **b**–**c,** Immunohistochemistry showing ProX-ID-dependent labelling in NAc, Thal, HT and CTX. **d**–**g,** Volcano plots for mPFC–NAc, mPFC–Thal, mPFC–HT and mPFC–CTX neural pathways. Dashed lines indicate FDR = 0.05 (horizontal) and log₂ fold change = −2 and 2 (vertical). Red dots represent significantly enriched proteins (FDR < 0.05, log_2_ fold change > 2). Gene symbols indicate the top 10 proteins with the largest fold changes. **h,** Heatmap of normalized log₂ biotinylation levels (ProX-ID vs. control) for representative presynaptic proteins. Asterisks indicate proteins significantly enriched in ProX-ID samples (FDR < 0.05, log_2_ fold change > 2, ProX-ID vs. control). NA indicates missing values. **i,** Odds ratios for selected Gene Ontology (GO) Cellular Component terms (see Materials and Methods). **j,** Proportion of known presynaptic, neurological disease-associated and functionally uncharacterized proteins among the neural pathway-enriched proteins. **k,** Venn diagram showing the overlap of significantly enriched proteins across mPFC–BLA, mPFC–NAc, mPFC–Thal, mPFC–HT and mPFC–CTX neural pathways. **l,** Top 20 enriched GO terms for Biological Process and Molecular Function.

GO Cellular Component annotation revealed predominant localization to synaptic vesicles and presynaptic active zones (Fig. 2i, Extended Data Fig. 4d–g). 60–80% of identified proteins overlapped with a previously reported presynaptic proteome (Fig. 2j, Extended Data Table 2), 1–2% of the proteins were functionally uncharacterized, and 30–40% were linked to neurological or psychiatric disorders (Fig. 2j, Extended Data Table 2 and 3). Comparative analyses revealed that 269 proteins were selectively enriched in mPFC–NAc, 12 in mPFC–Thal, 30 in mPFC–HT, two in mPFC–CTX and 174 in mPFC–BLA, whereas 77 proteins including Bsn and Syt1 were shared across all five pathways, representing a common presynaptic core (Fig. 2k, Extended Data Table 2). GO enrichment analysis showed that the five projection-specific proteome sets largely share the top 20 GO terms identified by Metascape ^44^ related to core synaptic communication (Fig. 2l, Extended Data Table 2). Despite shared categories, each pathway carried distinct molecular instances within annotated modules (Extended Data Fig. 5).

Protein–term networks based on the KEGG BRITE ^46–48^ database indicated pathway-dependent molecular composition across key functional domains (Fig. 3a, Supplemental Fig. 2–4). Integration with the KEGG database ^46–48^ visualized synaptic processes and highlighted pathway-specific profiles among intracellular signalling components, including Ca²⁺ channels (Slc8a1, Cacna1b, Cacna1c, Ryr2, Ryr3), adenylyl cyclases (Adcy1, Adcy2), protein phosphatases (Ppp3cb, Ppp1r37, Ppp3r1, Ppp1r9b, Ppp3ca, Ppp1r2), CaMKs and their associated proteins (Camk1d, Camk2d, Calm2), PKC-related proteins (Prkca, Prkcd, Frs2) and PKA-associated molecules (Prkar2a, Prkar2b, Csnk1e, Csnk1g3) (Fig. 3b). Proteins governing synaptic morphology also varied by neural pathway, spanning cytoskeletal components (Actg1), motor proteins (Dnal1, Kif2a, Myh6, Myo10), capping proteins (Cap1), the Rho GTPases (Rhob), Rho GAPs (Arhgap1, Arhgap23, Arhgap31, Arhgap39), Rho GEFs (Itsn1, Dock10, Dock11) and adhesion molecules (Sema4a, Alcam, Pecam1, Nectin1, Negr1, Lrrc4b, Ptprd, Slitrk3, Slitrk6, Nrxn1-α, Nrxn1-β, Cadm2, Cadm3, Cadm4, Cntn1, Nptn, Pcdh8, Igsf9b). The most prominent divergence occurred in vesicle trafficking machinery, with pathway-selective enrichment across endocytosis (Fnbp1, Fnbp1l, Lrp1b, Dab1, Wwp1, Hip1r, Cttn, Sh3glb2, Ap2s1, Eps15l1), exocytosis (Ppfia2), vesicular transporters (Slc17a6, Slc17a7), SNAREs (Stx7, Stxbp1, Stxbp5l, Stxbp6, Sec22b), endosome–lysosome transport proteins (Snx3, Vipas39), endosome–Golgi transport proteins (Clint1, Cog1, Sgasm1, Tmem165, Vps53) and Rab family proteins (Rab3a, Rab3c, Rab3gap2, Rab6a) (Fig. 3b). Neural pathway-specific protein–protein interaction networks are shown in Supplemental figures 5–7. To assess whether ProX-ID expression influenced detection depth, we ranked proteins by log_2_ fold change and compared the top 200 (Extended Data Fig. 6a, Extended Data Table 1, see Materials and Methods). Enrichment analysis consistently showed that while the same GO terms were enriched, the specific constituent proteins differed among projections (Extended Data Fig. 6b, Supplemental Fig. 8a–d). The protein–term network further showed that projection-selective proteins spanned diverse functional domains (Extended Data Fig. 6c). Together, these results demonstrate that mPFC neural pathways utilize distinct synaptic protein components, where conserved functions are implemented by neural pathway-specific molecular assemblies adapted to the functional demands of each target region.

**Figure 3.**
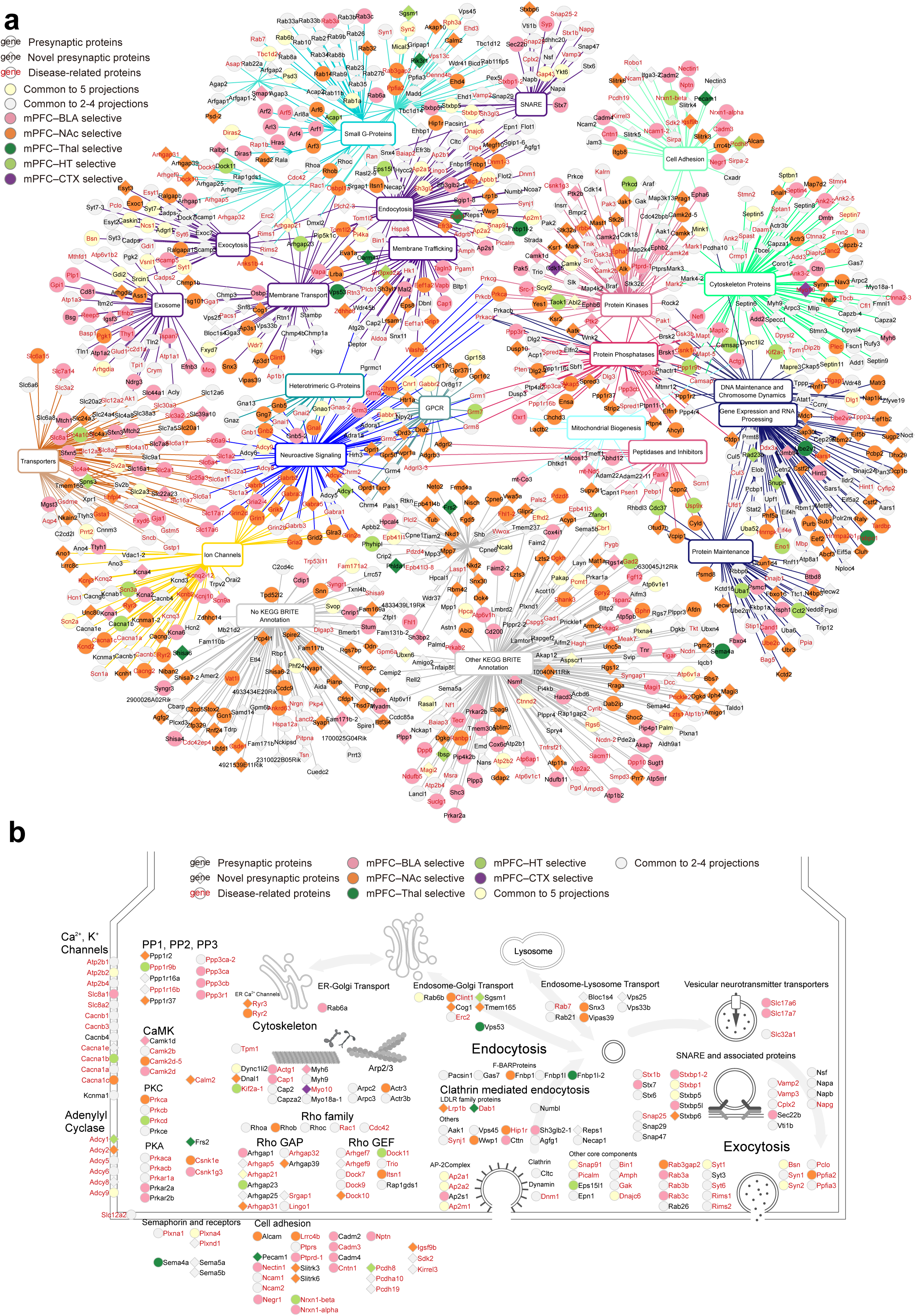
Spatial proteomics reveals molecular heterogeneity across mPFC projection pathways. **a,** KEGG BRITE-based protein–term network; nodes, proteins/terms; edges, associations based on BRITE functional categories. **b,** Projection-selective enriched proteins mapped onto representative presynaptic processes.

### BLTP2 is a novel presynaptic protein selectively enriched in the mPFC–BLA neural pathway

To further dissect the projection-enriched molecular composition, we prioritized proteins that were selectively enriched in the individual projections among the top 200 proteins with the largest log_2_ fold changes and that had not previously been reported as presynaptic or remained functionally uncharacterized (Fig. 4a, Extended Data Fig. 6). Of the two proteins exhibiting all three features, we focused on Bridge Like Lipid Transfer Protein Family Member 2 (BLTP2) for detailed molecular characterization within the mPFC–BLA pathway. In addition to its projection-enriched expression, recent human genetic studies have linked mutations in the BLTP2 gene to ASD and developmental delay, suggesting clinical relevance in neurodevelopmental disorders ^49–51^. Based on these findings, we investigated BLTP2 as a candidate molecule involved in projection-specific synaptic regulation and disease susceptibility. BLTP2 is a large protein comprising 2,235 amino acids, but its functional properties remain largely uncharacterized. Structural prediction using AlphaFold3 indicates that BLTP2 comprises an N-terminal transmembrane domain and a C-terminal region that adopts a tubule-like conformation (Fig. 4a).

**Figure 4.**
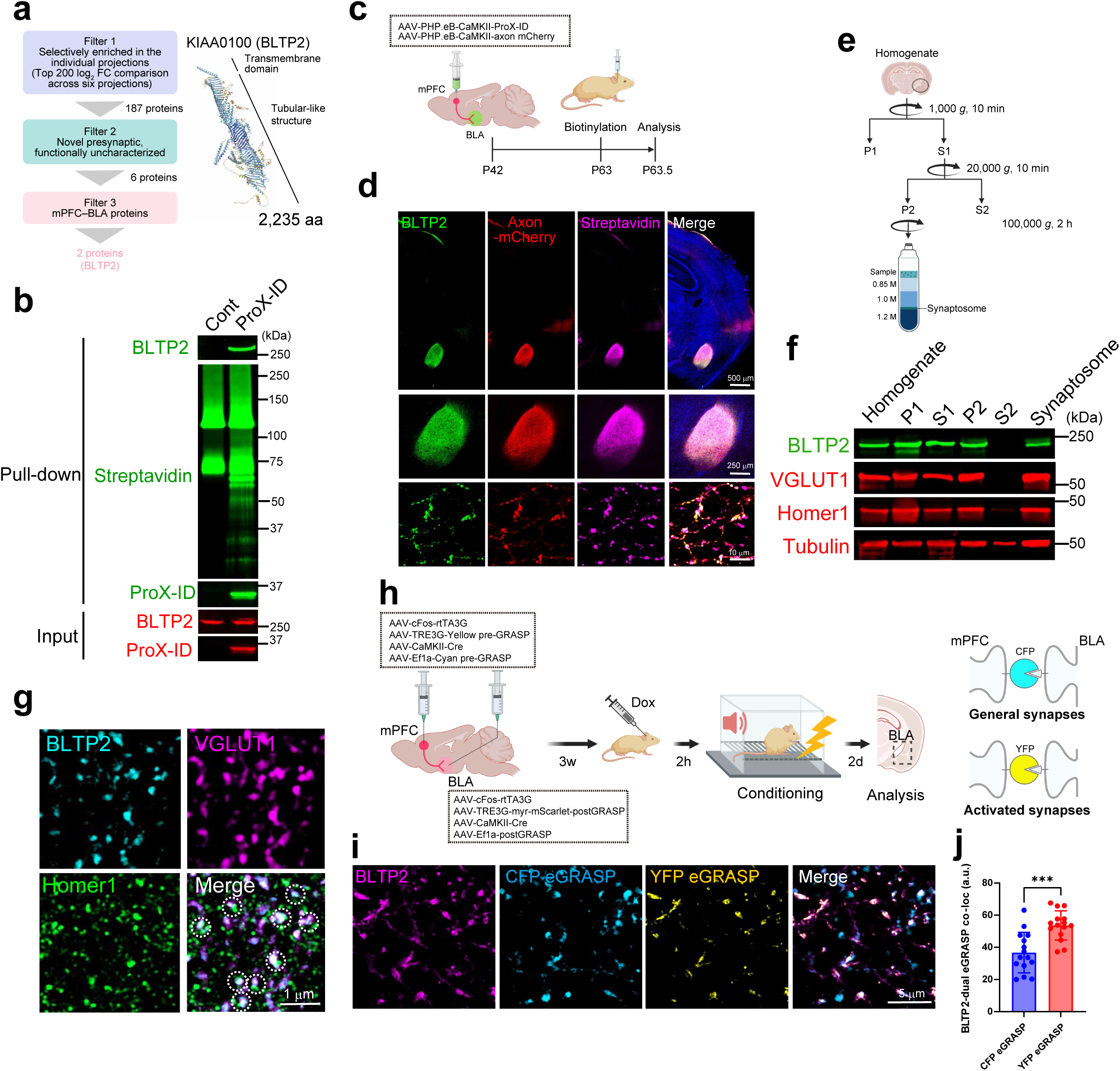
BLTP2 is a transmembrane synaptic protein that localizes to behaviourally activated mPFC–BLA synapses. **a,** Filtering process leading to the focus on BLTP2 (left). Schematic representation of the predicted structure of BLTP2 (right). **b,** Affinity purification followed by immunoblot analysis of biotinylated proteins from BLA tissue. **c,** Experimental design of *in vivo* ProX-ID labelling combined with axon-mCherry expression to visualize neural pathway-specific biotinylation and axonal projections from mPFC to BLA. **d,** Immunostaining of BLTP2 with axon-mCherry and biotinylation signals in the BLA. **e,** Experimental workflow of subcellular fractionation to isolate the synaptosomal fraction from BLA tissue. **f,** Immunoblot analysis of subcellular fractions from BLA lysates. **g,** Immunostaining of BLTP2 with excitatory synaptic markers Homer1 and VGLUT1 in the BLA. Dotted circles in the ‘Merge’ panel indicate colocalization of BLTP2 with Homer1- and VGLUT1-positive synapses. **h,** Dual-eGRASP strategy with BLTP2 immunostaining to visualize general (cyan eGRASP) and fear-activated (yellow eGRASP) mPFC–BLA synapses; AAV pre-/post-eGRASP constructs were injected into mPFC and BLA, with doxycycline given before conditioning. **i,** Representative images showing co-localization of BLTP2 with cyan and yellow eGRASP-labelled synapses in the BLA. **j,** Quantification of BLTP2 co-localization with CFP or YFP eGRASP-labelled synapses. Data are presented as mean ± s.e.m.; *n* = 15 slices per group from each of four mice. Student’s *t*-test. \*\*\**P* < 0.001.

To determine whether BLTP2 is biotinylated by ProX-ID selectively within the mPFC–BLA neural pathway, we isolated biotinylated proteins from BLA tissue of mice, which includes ProX-ID-expressing mPFC projection neurons targeting the BLA. We found that BLTP2 was biotinylated and enriched by ProX-ID within the mPFC–BLA neural pathway, whereas no significant enrichment was observed under control conditions (Fig. 4b). To further characterize the spatial distribution of BLTP2 in the mouse brain, we co-expressed axon-mCherry and ProX-ID in mPFC projection neurons to label their presynaptic terminals (Fig. 4c). Endogenous BLTP2 was prominently localized in the BLA, where it colocalized with both axon-mCherry and biotinylation signals, indicating its predominant presynaptic localization within the mPFC–BLA neural pathway (Fig. 4d). To determine whether BLTP2 is localized at synapses, we performed biochemical cell fractionation of BLA tissue (Fig. 4e). This revealed that BLTP2 is predominantly found in the membrane fraction (P2) rather than the cytoplasmic fraction (S2), consistent with its predicted domain structure featuring an N-terminal transmembrane domain (Fig. 4a and Fig. 4f). Moreover, BLTP2 was highly enriched in the synaptosomal fraction, where the excitatory synaptic markers VGLUT1 and Homer1 were also detected (Fig. 4f). Furthermore, immunohistochemical analysis showed that endogenous BLTP2 was highly colocalized with Homer1- and VGLUT1-positive excitatory synapses in the BLA (Fig. 4g).

Given the localization of BLTP2 at mPFC–BLA synapses, we next examined whether BLTP2 preferentially localizes to functionally active synapses by employing dual-eGRASP (dual-enhanced green fluorescent protein reconstitution across synaptic partners), a synapse-specific labelling technique that distinguishes between general anatomical synaptic connections and activity-dependent synaptic connections (Fig. 4h) ^52,53^. In this system, cyan eGRASP (CFP eGRASP) labels general synaptic contacts, whereas yellow eGRASP (YFP eGRASP) marks synapses that have been activated during specific behavioural experiences such as fear memory formation. To selectively label synapses within the mPFC–BLA neural pathway, we transduced AAV vectors carrying pre-eGRASP constructs (AAV-cFos-rtTA3G, AAV-TRE3G-Yellow pre-GRASP, AAV-CaMKII-Cre and AAV-Ef1a-Cyan pre-GRASP) into the mPFC, and post-eGRASP constructs (AAV-cFos-rtTA3G, AAV-TRE3G-myr-mScarlet-postGRASP, AAV-CaMKII-Cre and AAV-Ef1a-postGRASP) into the BLA (Fig. 4h). Doxycycline was administered prior to fear conditioning to induce activity-dependent expression of eGRASP components, enabling visualization of both general synaptic contacts and behaviourally activated synapses within the mPFC–BLA neural pathway (Fig. 4h). Quantitative analysis revealed that BLTP2 colocalized with approximately 36.8% of CFP eGRASP-labelled synapses, indicating its basal presence at anatomically defined mPFC–BLA synapses (Fig. 4i and j). Notably, BLTP2 exhibited a significantly higher colocalization rate (∼53.6%) with YFP eGRASP-labelled synapses, suggesting preferential enrichment at synapses activated during fear memory formation (Fig. 4i and j). Together, these findings demonstrate that ProX-ID enables projection-specific biotinylation of excitatory presynaptic proteins within the mPFC–BLA neural pathway and uncover BLTP2 as a previously uncharacterized synaptic protein that preferentially localizes to behaviourally activated synapses.

### BLTP2 is essential for the formation and functional maturation of excitatory synapses in the mPFC–BLA neural pathway

Given the specific localization of BLTP2 to excitatory presynaptic terminals within the mPFC–BLA neural pathway, we hypothesized that BLTP2 plays a pivotal role in the formation of excitatory synapses in this neural pathway. To test this, we first generated global BLTP2 knockout mice. However, homozygous deletion of BLTP2 resulted in embryonic lethality (data not shown), consistent with previous observations in *Drosophila* models lacking *KIAA0100*/*hobbit* ^54,55^. To circumvent this limitation, we employed a CRISPR/Cas9-based approach to generate projection-specific conditional knockout (cKO) mice, targeting the mPFC–BLA neural pathway. An AAV vector encoding three distinct single guide RNAs (sgRNAs) against BLTP2 (AAV-U6-sgRNA-BLTP2×3) was stereotaxically injected into the mPFC of Cas9-expressing mice (Fig. 5a), and efficient gene deletion in the targeted projection was confirmed by immunoblotting (Extended Data Fig. 7a and b). Quantitative analysis revealed that BLTP2-cKO mice exhibited a significant reduction in excitatory synapse density within the BLA as determined by the co-localization of Homer1 with VGLUT1 (Fig. 5b and c). To validate this phenotype using an independent approach, we employed two shRNAs targeting BLTP2 (shBLTP2#1 and shBLTP2#2), which efficiently reduced BLTP2 protein level by >80% in cultured neurons (Extended Data Fig. 7c and d). Knockdown of BLTP2 significantly decreased the density of excitatory synapses both *in vitro* (Extended Data Fig.7e and f) and *in vivo* (Extended Data Fig.7g and h), indicating the robustness and reproducibility of the phenotype across both experimental systems. To examine whether BLTP2 contributes to presynaptic structural organization, we assessed the volume of presynaptic boutons. Morphometric analyses revealed a significant reduction in presynaptic bouton volume in the BLA of BLTP2-cKO mice compared to controls (Fig. 5d). Consistent with these findings, neural pathway-specific BLTP2 knockdown also led to decreased presynaptic bouton size (Extended Data Fig.7i).

**Figure 5.**
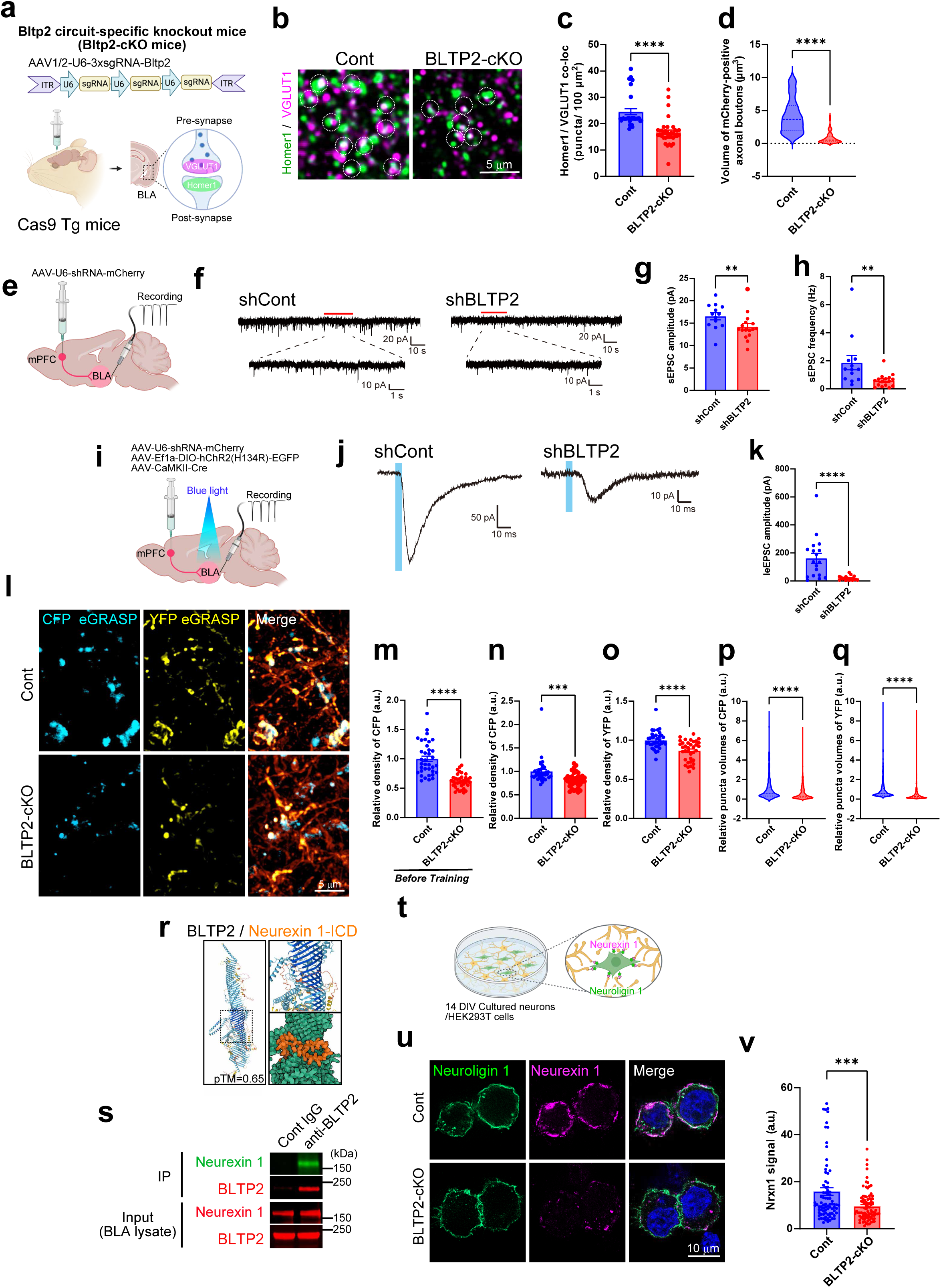
BLTP2 controls synapse formation, transmission and activity-dependent remodelling in the mPFC–BLA pathway. **a,** Projection-specific BLTP2 knockout in mPFC–BLA using AAV-U6-sgRNA in Cas9 transgenic mice; BLA sections were analysed for Homer1 and VGLUT1. **b,** Representative images of BLA sections immunostained for Homer1 (green) with VGLUT1 (magenta; upper panels) in control and BLTP2-cKO mice. **c,** Quantification of the synaptic density of Homer1 and VGLUT1 double-positive excitatory synapses in the BLA. Data are presented as mean ± s.e.m.; *n* = 36 slices per group from each of four mice. Student’s *t*-test. \*\*\*\**P* < 0.0001. **d,** Presynaptic bouton volume distributions (violin plots); bold dashed line, median; thin dashed lines, quartiles; *n* = 100 slices per group from each of four mice. Student’s *t*-test. \*\*\*\**P* < 0.0001. **e,** Schematic illustration of whole-cell patch-clamp recordings from BLA pyramidal neurons. **f,** sEPSC traces after projection-specific expression of shCont or shBLTP2. **g**–**h,** Quantification of sEPSC amplitude (**g**) and frequency (**h**). Data are presented as mean ± s.e.m.; shCont *n* = 13, shBLTP2 *n* = 16 cells from each of three mice. Two-tailed Mann–Whitney test. \*\**P* < 0.01. **i,** Optogenetic stimulation and recordings from BLA pyramidal neurons expressing hChR2 (H134R). **j,** leEPSCs recorded from BLA pyramidal neurons following mPFC–BLA neural pathway-specific expression of shCont or shBLTP2. **k,** Quantification of leEPSC amplitude. Data are presented as mean ± s.e.m.; shCont *n* = 18, shBLTP2 *n* = 17 cells from each of three mice. Two-tailed Mann–Whitney test. \*\*\*\**P* < 0.0001. **l,** CFP (cyan) and YFP (yellow) eGRASP-labelled synapses in the BLA following fear conditioning. **m**–**o,** Quantification of the synaptic density of CFP eGRASP-labelled basal synapses and YFP eGRASP-labelled learning-induced synapses after fear conditioning. Data are presented as mean ± s.e.m.; *n* = 36 slices per group from each of four mice. Student’s *t*-test. \*\*\*\**P* < 0.0001; \*\*\**P* < 0.001. **p**–**q,** Quantification of the synaptic volume of CFP eGRASP-labelled basal synapses and YFP eGRASP-labelled learning-induced synapses after fear conditioning. Cont *n* = 650 (CFP) and 746 (YFP), BLTP2-cKO *n* = 612 (CFP) and 891 (YFP) synapses from each of four mice. Student’s *t*-test. \*\*\*\**P* < 0.0001. **r,** Structural model of BLTP2 and the intracellular domain of Neurexin 1, suggesting potential interaction sites predicted by pTM score. **s,** Co-immunoprecipitation showing physical interaction between BLTP2 and Neurexin 1 from BLA lysates. **t,** Schematic of co-culture assay. **u,** Representative images showing subcellular localization of Neuroligin 1 (green) and Neurexin 1 (magenta) in BLTP2 knockout neurons and control. **v,** Quantification of Neurexin 1 (Nrxn1) signal intensity in Neuroligin 1-positive regions. Data are presented as mean ± s.e.m.; Cont *n* = 82, BLTP2-cKO *n* = 85 neurons. Student’s *t*-test. \*\*\**P* < 0.001.

We next investigated whether BLTP2 deficiency affects synaptic function. Whole-cell patch-clamp recordings from BLA pyramidal neurons in mice with neural pathway-specific BLTP2 knockdown revealed a significant decrease in both the frequency and amplitude of spontaneous excitatory postsynaptic currents (sEPSCs) compared to controls (Fig. 5e–h). Furthermore, optogenetic activation of mPFC–BLA projections using humanized Channelrhodopsin-2 H134R (hChR2(H134R)) elicited robust light-evoked EPSCs (leEPSCs) in control mice, whereas BLTP2 knockdown markedly attenuated leEPSC amplitudes (Fig. 5i–k), indicating impaired synaptic transmission efficacy.

To evaluate the role of BLTP2 in activity-dependent synaptic remodelling, we employed the dual-eGRASP system to visualize synaptic changes in the mPFC–BLA neural pathway following fear conditioning. In BLTP2-cKO mice, both CFP- and YFP-eGRASP-labelled synapses were significantly reduced in number and volume compared to controls after conditioning (Fig. 5l–q), indicating that BLTP2 is essential for the experience-dependent formation and maturation of excitatory synapses. Notably, even under baseline conditions (prior to conditioning), BLTP2-cKO mice exhibited a significant reduction in CFP-eGRASP-labelled synapses in the mPFC–BLA neural pathway (Fig. 5m), suggesting that BLTP2 is also required for maintaining basal synaptic connectivity. Consistent with these results, shBLTP2-expressing neurons in the mPFC–BLA neural pathway displayed a marked reduction in CFP-eGRASP-labelled synapses under baseline conditions (Extended Data Fig.7j–l), indicating selective involvement of BLTP2 in maintaining basal synaptic connectivity. In addition, following fear conditioning, both CFP- and YFP-eGRASP-positive puncta were significantly reduced in number and volume in shBLTP2-expressing neurons in the mPFC–BLA neural pathway (Extended Data Fig. 7m–p), demonstrating that BLTP2 is also critical for activity-dependent synaptic plasticity.

Next, we examined the molecular mechanism underlying BLTP2-mediated synapse formation in the mPFC–BLA neural pathway. We hypothesized that BLTP2 interacts with Neurexin 1, a pivotal presynaptic organizer involved in the assembly of diverse synaptic structures ^3^. Structural prediction using AlphaFold3 revealed a high-confidence interaction between BLTP2 and the intracellular domain of Neurexin 1 (pTM = 0.65; Fig. 5r). Co-immunoprecipitation from BLA lysates confirmed the interaction of BLTP2 with Neurexin 1 *in vivo* (Fig. 5s). To determine whether BLTP2 contributes to synapse formation via Neurexin 1–Neuroligin 1 interactions, we utilized a co-culture system in which primary cortical neurons were cultured together with HEK293T cells expressing Neuroligin 1 (Fig. 5t). In this assay, control neurons exhibited robust accumulation of Neurexin 1 at sites of contact with Neuroligin 1-expressing HEK293T cells, reflecting the formation of trans-synaptic adhesive complexes (Fig. 5u and v). In contrast, BLTP2-deficient neurons failed to accumulate Neurexin 1 at these contact sites (Fig. 5u and v). This deficit was also recapitulated by shRNA-mediated knockdown of BLTP2, indicating that BLTP2 is essential for the presynaptic recruitment of Neurexin 1 and the formation of stable Neurexin–Neuroligin interactions (Extended Data Fig. 7q and r). Collectively, these findings demonstrate that BLTP2 plays an essential role in both the structural and functional establishment of excitatory synapses in the mPFC–BLA neural pathway by regulating presynaptic Neurexin 1 localization.

### Loss of BLTP2 in the mPFC–BLA pathway impairs contextual fear memory, reduces anxiety, and disrupts social interaction

Recent genomic studies have implicated the BLTP2 gene in multiple neuropsychiatric and neurodevelopmental disorders including ASD, schizophrenia and developmental delay ^49–51^. Notably, the mPFC–BLA neural pathway has been functionally linked to emotional learning, anxiety-related behaviour and social interaction ^21,22,56,57^. To investigate whether loss of BLTP2 in the mPFC–BLA neural pathway alters these behaviours, we assessed BLTP2-cKO mice with the open field test, elevated plus maze (EPM), three-chamber social interaction test and fear conditioning test (Fig. 6). In auditory fear conditioning tests, BLTP2-cKO mice showed freezing responses comparable to controls in conditioning trials and auditory cues, but a significantly reduced freezing response in the contextual test (Fig. 6a–e), consistent with reduced activity-dependent synapse formation after conditioning (Fig. 5l, o, q). Thus, BLTP2 is required for contextual fear-memory formation, but not for fear-memory acquisition or tone-cued retrieval, aligning with evidence that contextual and auditory fear memories rely on hippocampus–mPFC–BLA and auditory cortex–BLA pathways, respectively ^56,57^. In the open field test, BLTP2-cKO mice showed no significant differences from control mice in total locomotor activity or time spent in the centre zone (Fig. 6f–h), indicating normal exploratory behaviour and spatial anxiety. On the other hand, BLTP2-cKO mice spent significantly more time in the open arms of the EPM compared to controls (Fig. 6i–k), indicating reduced anxiety-like behaviour. In the three-chamber social interaction test ^58,59^, control mice exhibited a strong preference for the chamber containing an unfamiliar conspecific (i.e., social stimulus) over an empty cage, whereas BLTP2-cKO mice displayed a significantly reduced preference for the social stimulus (Fig. 6l–o), suggesting impaired social interaction. Taken together, these findings demonstrate that BLTP2 selectively regulates contextual fear memory formation, anxiety-like behaviour and social interaction. The convergence of these phenotypes with those reported in established ASD mouse models supports a role for BLTP2 in mPFC–BLA circuit dysfunction underlying emotional and social behavioural deficits ^22,60–64^.

**Figure 6.**
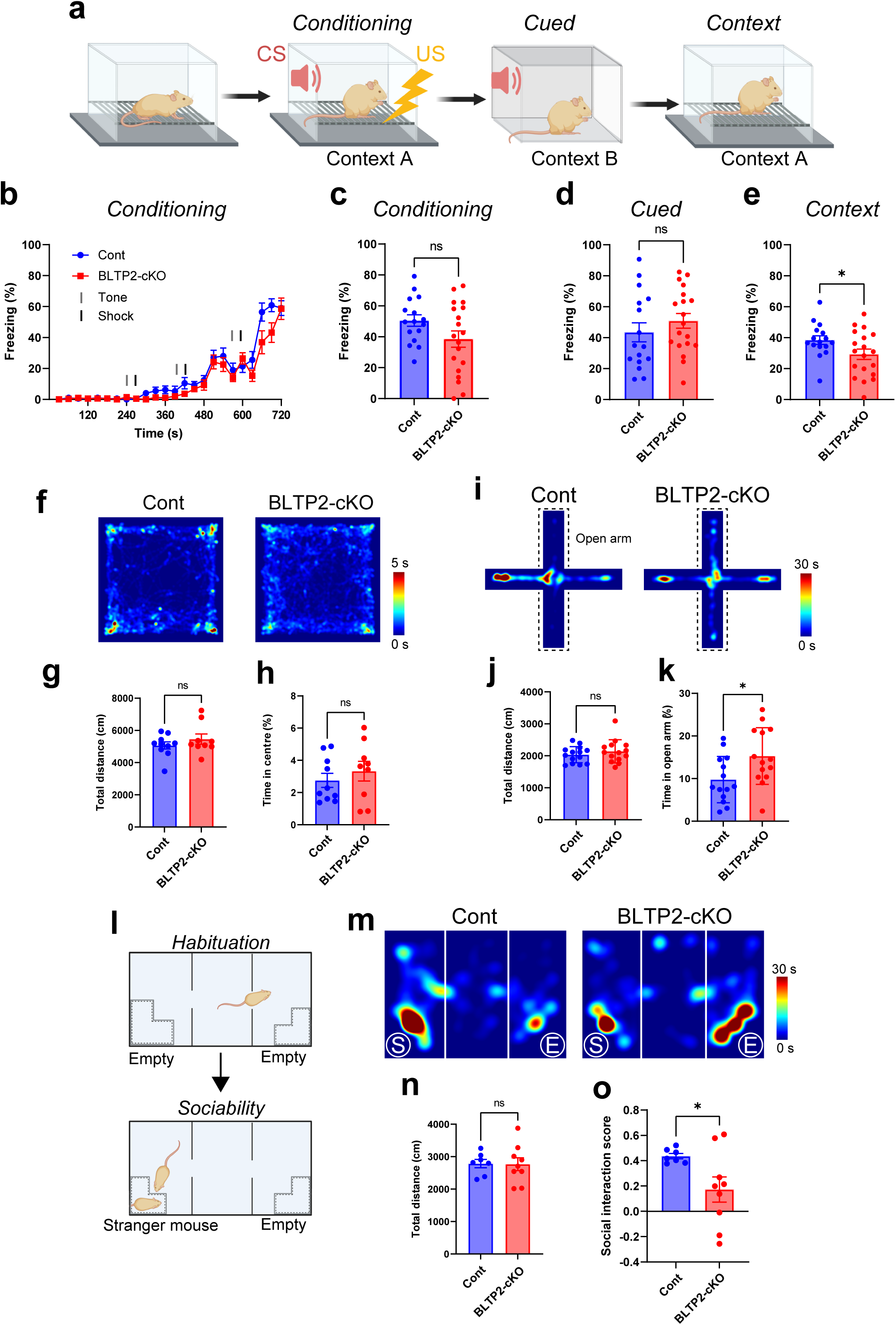
Loss of BLTP2 in the mPFC–BLA pathway disrupts fear memory, anxiety regulation and social behaviour. **a,** Schematic of the fear conditioning protocol. **b,** Percentage of freezing duration during conditioning. Data are presented as mean ± s.e.m.; Cont *n* = 16, BLTP2-cKO *n* = 19 mice. **c**–**e,** Freezing during the post-tone in the fear conditioning trial (**c**), cued trial (**d**) and contextual trial (**e**). Data are presented as mean ± s.e.m.; Cont *n* = 16, BLTP2-cKO *n* = 19 mice. Student’s *t*-test. \**P* < 0.05; ns, not significant. **f,** Heatmaps showing the time distribution during the open field test. **g**–**h,** Quantification of total distance travelled (**g**) and percentage of time spent in the centre zone (**h**). Data are presented as mean ± s.e.m.; Cont *n* = 10, BLTP2-cKO *n* = 9 mice. Student’s *t*-test. ns, not significant. **i,** Heatmaps showing the time distribution during the elevated plus maze (EPM) test. **j**–**k,** Quantification of total distance travelled (**j**) and percentage of time spent in the open arms (**k**) in EPM. Data are presented as mean ± s.e.m.; Cont *n* = 14, BLTP2-cKO *n* = 14 mice. Student’s *t*-test. \**P* < 0.05; ns, not significant. **l,** Schematic of the three-chamber social interaction test. **m,** Heatmaps showing the time distribution during the three-chamber interaction test. **n,** Quantification of total distance travelled. **o,** Social interaction score. Data are presented as mean ± s.e.m.; Cont *n* = 7, BLTP2-cKO *n* = 9 mice. Student’s *t*-test. \**P* < 0.05; ns, not significant.

## Discussion

Synapses exhibit remarkable molecular diversity, but how this diversity is specialized across anatomically defined neural circuits remains largely unexplored. Our comprehensive spatial proteomic analyses using the ProX-ID approach revealed that each mPFC-projecting neural pathway exhibits a distinct presynaptic proteomic architecture, characterized by pathway-specific enrichment of proteins involved in vesicle trafficking, cytoskeletal organization, synaptic adhesion and intracellular signaling (Fig. 3, Extended Data Fig. 6). Among these core synaptic functions, the molecular constituents were highly pathway-selective (Fig. 3). For example, proteins related to synaptic vesicle trafficking (e.g., Eps15l1, Stx7, Slc17a6, Slc17a7), cytoskeletal dynamics (e.g., Actg1, Kif2a, Rho GTPases), synaptic adhesion (e.g., Nrxn1, Nectin1, Slitrk3) and signal transduction (e.g., Adcy1, PP1 family, PKC-related proteins) displayed projection-specific enrichment, indicating that individual pathways deploy distinct molecular components to implement shared synaptic processes (Fig. 3). Cross-species analysis showed that most proteins enriched in the mouse mPFC–BLA pathway had orthologues that are expressed in the marmoset prefrontal cortex (Fig. 1k), and many of these conserved proteins are associated with human neuropsychiatric and neurodevelopmental disorders (Extended Data Fig. 3b). Because mPFC-centred circuits are deeply involved in fear, anxiety and social behaviours and have been repeatedly implicated in disorders such as ASD, schizophrenia, post-traumatic stress disorder and major depression, future integration of the ProX-ID atlas with other public datasets such as genome-wide association studies and large-scale gene expression and regulatory data from human postmortem brain tissues (PsychENCODE ^65,66^) should provide a reusable resource for overlaying known circuit functions and risk-associated genes onto projection-resolved presynaptic modules in cases where behavioural roles or genetic associations are known but synaptic molecular mechanisms remain unclear.

This molecular diversity likely underlies the functional specialization observed across distinct neural circuits. While previous studies have focused on candidate molecules within individual circuits, increasing evidence indicates that synaptic protein expression varies across projections and is critical for specifying, maturing and modulating synaptic connections. Electrophysiological recordings have shown that circuits involving the mPFC, BLA and NAc operate at distinct frequency bands, which likely reflects differences in synaptic properties shaped by molecular composition ^67^. Behavioural experiments further demonstrate that these anatomically segregated circuits serve distinct functions ^68–70^. In particular, our findings demonstrate that BLTP2-controlled mPFC–BLA pathway contributes to fear memory formation, anxiety-like behaviour and social novelty preference (Fig. 6). These results are consistent with previous reports implicating this projection in broader functions such as emotional memory processing, threat evaluation and social behaviour^21,22,27,28,56,57,69,71^. These observations suggest that even within anatomically defined pathways, molecularly distinct neuronal subpopulations can support specialized behavioural functions. Our spatially resolved proteomic approach enables systematic investigation of these projection-specific molecular signatures and provides a foundation to link molecular diversity with electrophysiological properties and behavioural output of neural circuits.

Despite major efforts to characterize postsynaptic compartments using biochemical fractionation and affinity purification, the presynaptic proteome remains underexplored, largely due to the structural complexity of presynaptic terminals and the lack of labelling strategies compatible with intact anatomical context. Recent methods such as fluorescence-activated synaptosome sorting (FASS) and peroxidase-based proximity labelling (e.g., HRP and APEX) have advanced our understanding of cell type-specific synaptic diversity, but they rely on tissue homogenization or potentially cytotoxic reagents and do not readily resolve projection-defined synapses *in vivo* ^18,72,73^. FASS disrupts cytoarchitecture and reduces confidence in assigning pre- versus postsynaptic origin, whereas HRP and APEX require hydrogen peroxide, which limits their compatibility with live tissue and long-term behavioural paradigms ^33^. ProX-ID addresses this methodological gap by selectively labelling presynaptic terminals in anatomically defined projections under near-physiological conditions, without exogenous hydrogen peroxide, synaptosome purification or overexpression of large synaptic tags, thereby preserving neuronal viability and anatomical integrity (Fig. 1–3, Extended Data Fig. 8). Using this workflow, presynaptic proteomes can be directly compared across multiple projection pathways within the same brain, revealing pathway-specific molecular profiles and potential vulnerabilities relevant to neuropsychiatric disease.

Among the pathway-selective proteins identified, we focused on BLTP2 (KIAA0100), a previously uncharacterized transmembrane protein that is highly enriched in the mPFC–BLA circuit. BLTP2 is evolutionarily conserved, and its *Drosophila* orthologue *hobbit* is essential for regulated exocytosis and viability ^54^. Consistent with this, BLTP2 knockout mice exhibited embryonic lethality (data not shown). Although recent work implicates BLTP2 in organizing ER–plasma membrane contact sites via FAM102 family proteins ^74^, its neural function has remained unclear. Here, we show that BLTP2 is selectively localized to excitatory presynaptic terminals in the mPFC–BLA pathway and becomes further enriched at synapses that are activated during fear memory formation (Fig. 4). Loss of BLTP2 in this projection selectively impaired contextual fear memory retrieval, reduced anxiety-like behaviour, and diminished social interaction, without affecting general locomotion (Fig. 6). These behavioural phenotypes coincided with presynaptic defects, smaller boutons, weakened excitatory transmission and blunted activity-dependent remodelling (Fig. 5). Mechanistically, BLTP2 recruits Neurexin 1 to stabilize projection-specific synaptic integrity (Fig. 5). Human genetics supports disease relevance, with rare *de novo* BLTP2 mutations reported in individuals with ASD, schizophrenia and intellectual disability ^49–51^. BLTP2 is also highly expressed in socioemotional brain regions such as the BLA, and BLTP2-deficient mice recapitulate behavioural features observed in rodent models of social anxiety, including reduced social approach and increased avoidance ^64^. These findings suggest that BLTP2 functions as a projection-specific synaptic organizer that integrates emotional and social information, and that its dysfunction confers selective vulnerability in circuits implicated in neuropsychiatric disorders.

In conclusion, our work provides a framework for understanding how molecular diversity at presynaptic terminals underlies pathway specific synaptic specialization and behavioural function, and establishes ProX-ID as a broadly accessible platform for *in vivo* projection specific presynaptic proteomics (Extended Data Fig. 8). By integrating projection resolved spatial proteomics with functional and behavioural analyses, we show that long range projections deploy distinct molecular networks to shape synaptic connectivity and regulate behaviour. Together, this mPFC centred presynaptic atlas provides a foundation for investigating synaptic vulnerabilities in complex brain disorders and for developing circuit targeted molecular interventions guided by pathway specific presynaptic architectures.

## Acknowledgements

We thank Prof. Scott H. Soderling (Duke University) for providing valuable suggestions on the experimental design of ProX-ID. We thank Dr. Shota Morikawa (Tokyo University) for providing valuable suggestions on the experimental design of behavioural analysis. We would like to thank Dr. Ian Smith for English language editing. This work was supported by a Grant-in-Aid for Scientific Research B (21380936), PRESTO (21461219 and 24029397) from JST (T.T.), a Brain Mind 2.0 from AMED (JP23wm0625001 for T.S. and 24019272, 24019528 for T.T.), The Ono Pharmaceutical Foundation for Oncology (T.T., T.M.), Immunology and Neurology (T.T., T.M.), the Memorial Foundation for Medical and Pharmaceutical Research (T.T.), a Research Grant from the JGC Saneyoshi Scholarship Foundation (T.T.), the Takeda Science Foundation (T.T., T.M.), MEXT Grants-in-Aid for Scientific Research on Innovative Areas “Cell type census of adaptive neuronal circuits: biological mechanisms of structural and functional organization (Adaptive Circuit Census, ACC)” (24H01250) (T.T.). This work was also supported by Joint Usage and Joint Research Programs of the Institute of Advanced Medical Sciences, Tokushima University (T.T.). This work was also supported by JSPS KAKENHI Grants (JP20H05900 for M.T. and JP24H00067 for M.T.) and the Core Research for Evolutional Science and Technology (CREST) program from AMED under Grant Number 24gm1510013h (M.T.). S.U. was a JSPS research fellow (JP23KJ1729). This work was also supported by the MEXT Cooperative Research Project Program, Medical Research Center Initiative for High Depth Omics, and CURE:JPMXP1323015486 for MIB, and AMRC, Kyushu University, and by AMED (JP23gm1910004, JP23jf0126004, JP24zf0127012 for T.M.), JSPS KAKENHI (JP22H05062, JP25H01009, JP25K02573 for T.M.), The Mitsubishi Foundation (T.M.), Daiichi Sankyo Foundation of Life Science (T.M.), Mochida Memorial Foundation for Medical and Pharmaceutical Research (T.M.), Astellas Foundation for Research on Metabolic disorders (T.M.), The Nakajima Foundation (T.M.) and The Uehara Memorial Foundation (T.M.). Some figure elements were created with BioRender.com.

## Author contribution

Y.I. and T.T. designed the study. Y.I., S.U., M.T., H.K. and T.T. wrote the manuscript. S.N., M.F. and T.T. produced the constructs. Y.I., S.N., H.K. and T.T. performed *in vivo* BioID-based proteomics analysis. K.O. and T.S. performed spatial transcriptomic analysis in mouse and marmoset brains. S.N., J.M. and T.T. performed the biological experiments. S.N., J.M., M.F. and T.T. performed imaging and morphological analyses. S.U., Y.K. and M.T. performed electrophysiological analysis. Y.I., A.F., S.H., K.T., M.K., T.M., Y.H. and T.T. performed the behavioural analysis. All authors discussed the results and commented on the manuscript text.

## Data availability

Data supporting the figures will be made available upon peer-reviewed publication.

## Methods

### Animals

Wild-type C57BL/6J mice were purchased from KYUDO Company (Breeder: Jackson Laboratory Japan). All mice were housed in the specific pathogen-free animal facility at Kyushu University with a 12-h light/dark cycle. Only male mice were used for behavioural testing, and both sexes were used for other experiments. All experimental procedures in mice were approved by the Kyushu University Animal Experiment Committee (A24-335-1), and the care of the animals was in accordance with institutional guidelines. All experiments were performed to minimize animal pain, stress and the number of animals.

### Plasmid construction

pSF3-ultraID was a gift from Julien Béthune (Addgene plasmid #172878), and pAAV-hSyn-Flex-Axon-EGFP was a gift from Rylan Larsen (Addgene plasmid #135429). ProX-ID was subcloned with ultraID and the GAP43 palmitoylation sequence into the pAAV-CaMKII vector. Axon-mCherry was subcloned into the pAAV-CaMKII vector. Ef1a-DIO-mCherry was amplified and subcloned into AAV-U6-shRNA vectors. The shRNA sequences used were as follows: shCont, 5′-GATCACGGATTCGAGAACAGA-3′; shBLTP2#1, 5′-GGATCCAGAATGTCAGTCTT-3′; shBLTP2#2, 5′-GCCATGATCTGCCAAACTAT-3′. pAAV-TIWB-myrmScarlet-I-P2A-post-eGRASP (Addgene plasmid #111584), pAAV-EWB-DIO-myriRFP670V5-P2A-post-eGRASP (Addgene plasmid #111585), pAAV-EWB-DIO-cyan pre-eGRASP(p32) (Addgene plasmid #111589), pAAV-TIWB-yellow pre-eGRASP(p30) (Addgene plasmid #111588) and pAAV-FAH-rtTA3G (Addgene plasmid #120309) were gifts from Bong-Kiun Kaang. pCAG-HA-Nrxn1alpha was a gift from Peter Scheiffele (Addgene plasmid #58266). NGL-1-myc/pEGFP-N1 was a gift from Eunjoon Kim (Addgene plasmid # 197314). pAAV-EF1a-DIO-hChR2(H134R)-P2A-EYFP was a gift from Mingshan Xue (Addgene plasmid #139283). pAAV-pMecp2-SpCas9-spA (PX551) (Addgene plasmid #60957) and PX552 (pAAV-U-6sgRNA (SapI)_hSyn-GFP-KASH-bGH) (Addgene plasmid #60958) were gifts from Feng Zhang. The sgRNA sequences used were as follows: sgRNA-Scrambles, 5′-GCGATCCCATGCCCTTAGAA-3′, 5′-GCGACCTCCCACGCAAGAAA-3′, 5′-GGCGCAACCGGCAATAGAGA-3′; sgRNA-BLTP2, 5′-GACTCCACTGAGACTGATCCG-3′, 5′-GCAAAGCCAGCAGCCAACTCG-3′, 5′-GCATAGAGACGACGAGAGCGC-3′. All constructs were confirmed by DNA sequencing. All primers are shown in Supplemental Table 1.

### Antibodies

The following antibodies were used: rabbit anti-HA (Cell Signaling Technology, C29F4), rabbit anti-Homer1 (Synaptic Systems, 160-003), guinea pig anti-VGLUT-1 (Synaptic Systems, 135-304), monoclonal α-Tubulin (Santa Cruz, sc-32293), rabbit anti-KIAA0100 (BLTP2) (MyBioSource, MBS8243055), rabbit anti-mCherry (Abcam, ab167453), rabbit anti-Neurexin-1α (Nittobo Medical, MSFR104630), chicken anti-GFP (Merck, AB16901), Alexa Fluor 488 Goat anti-Mouse (ThermoFisher, A32723), Alexa Fluor 488 Goat anti-Rabbit (ThermoFisher, A-11008), Alexa Fluor 488 Goat anti-Guinea pig (ThermoFisher, A11073), Alexa Fluor 488 Goat anti-Chicken (ThermoFisher, A-11039), Alexa Fluor 555 Goat anti-Rabbit (ThermoFisher, A21428), Alexa Fluor 555 Goat anti-Guinea pig (ThermoFisher, A21435), Alexa Fluor 647 Donkey anti-rabbit (ThermoFisher, A31573), Alexa Fluor 647 Goat anti-Chicken (ThermoFisher, A-21449), Alexa Fluor 647 Donkey anti-Guinea pig (Jackson ImmunoResearch, 706-605-148), Alexa Fluor 647 Streptavidin (ThermoFisher, S21374), StarBright Blue 700 anti-Mouse (BioRad, 12004159), StarBright Blue 700 anti-Rabbit (BioRad, 12004162), CF770 anti-Mouse (Biotium, 20077), CF770 anti-Rabbit (Biotium, 20078) and DAPI solution (Dojindo, 340-07971).

### AAV production

AAVs were produced as previously described ^39^. Briefly, HEK293T cells were transfected with pAd-DELTA F6, serotype plasmid AAV PHP.eB or AAV 1/2 and AAV plasmid. After 72 h, the cells were lysed in 15 mM NaCl, 5 mM Tris-HCl, pH 8.5, and incubated with 50 U/ml Benzonase for 30 min at 37°C. The cell extracts including AAVs, which were obtained by centrifugation at 2,800 × *g* for 30 min at 4°C, were centrifuged on a gradient of 15%, 25%, 40% and 60% iodixanol solution using a Beckman Ti-70 rotor, spun at 67,000 rpm for 1 h. Virus particles were recovered from the boundary between 40% and 60% iodixanol and concentrated with a 100-kDa filter (Vivaspin, NIPPON Genetics). Virus titers were measured by qPCR using a linearized genome plasmid as a standard (∼2 × 10^13^ genome copies /mL) ^39^. For small-scale AAV supernatant, HEK293T cells were transfected with pAd-DELTA F6, serotype plasmid AAV PHP.eB or AAV 1/2 and AAV plasmid. After 72 h, the AAV-containing supernatant medium was collected and filtered with a 0.45-μm cellulose acetate Spin-X centrifuge tube filter (Costar 8162).

### Primary neuronal culture and HEK293T cell cultures

Cortical neurons were prepared from E15 mouse embryos with papain as previously described ^39,75^. These cells were seeded on coverslips or dishes coated with poly-D-lysine (Sigma) and cultured in neurobasal medium (Invitrogen) supplemented with B-27 (Invitrogen) and 1 mM GlutaMAX (Invitrogen). HEK293T (obtained from ATCC; #CRL-11268) cells were maintained in DMEM (Gibco) supplemented with 10% FBS (Gibco) and 100 U/mL penicillin/streptomycin. Cell lines were incubated at 37°C in 5% CO_2_. Cells were passaged every three days.

### Stereotaxic surgery

Mice were deeply anaesthetized via an intraperitoneal injection of 0.1 M phosphate-buffered saline (PBS) containing a three-agent anaesthetic mixture composed of medetomidine (0.75 mg/kg), midazolam (4 mg/kg) and butorphanol (5 mg/kg) at standard concentrations. Once anaesthetized, each mouse was secured in a stereotaxic apparatus (Narishige), and ophthalmic ointment was applied to protect the eyes. After removing scalp hair and disinfecting the skin with Betadine, a midline scalp incision was made to expose the skull. Craniotomies were then performed using a microdrill at the designated coordinates for viral injections. Viral injections were performed using a glass capillary (World Precision Instruments, 504950) connected to a Nanoliter 2020 injector (World Precision Instruments). Vectors were injected at a rate of 85 nL/min, and the needle was left in place for an additional 1 min to allow for diffusion before being withdrawn. Following injection, the scalp was closed using surgical sutures and tissue adhesive. Post-surgery, mice were placed in a recovery cage under a heat lamp until they regained consciousness, and were allowed at least two weeks to recuperate prior to further procedures or behavioural testing.

For *in vivo* BioID experiments (Fig. 1, Fig. 2, Extended Data Fig. 1 and Extended Data Fig. 4), C57BL/6J mice received bilateral injections of 800 nL AAV-PHP.eB-CaMKII-ProX-ID into either the mPFC or the BLA. For labelling mPFC–BLA projection neurons (Extended Data Fig. 1g and Fig. 4d), C57BL/6J mice received bilateral injections of 800 nL AAV-CaMKII-axon-mCherry into the mPFC. For labelling mPFC–BLA synapses and activated synapses using dual-eGRASP (Fig. 4i, 5l and Extended Data Fig. 7k), C57BL/6J mice received bilateral injections into the mPFC of a total volume of 1,200 nL containing AAV-PHP.eB-CaMKII-Cre, AAV-PHP.eB-FAH-rtTA3, AAV-PHP.eB-EWB-DIO-cyan pre-eGRASP (p32) and AAV-PHP.eB-TIWB-yellow pre-eGRASP (p30). In addition, a total volume of 1,200 nL containing AAV-PHP.eB-CaMKII-Cre, AAV-PHP.eB-FAH-rtTA3, AAV-PHP.eB-TIWB-myrmScarlet-I-P2A-post-eGRASP and AAV-PHP.eB-EWB-DIO-myriRFP670V5-P2A-post-eGRASP was injected bilaterally into the BLA.

For projection-specific knockdown experiments (Extended Data Fig. 7g–r), C57BL/6J mice received bilateral mPFC injections of a total volume of 800 nL containing AAV-PHP.eB-CaMKII-Cre together with either AAV-PHP.eB-EF1a-DIO-mCherry-U6-shCont, AAV-PHP.eB-EF1a-DIO-mCherry-U6-shBLTP2#1 or AAV-PHP.eB-EF1a-DIO-mCherry-U6-shBLTP2#2. For additional knockdown experiments using SpCas9, C57BL/6J mice received bilateral mPFC injections of a total volume of 800 nL containing pAAV-pMecp2-SpCas9-spA together with either AAV-PHP.eB-EF1a-DIO-mCherry-U6-shCont, AAV-PHP.eB-EF1a-DIO-mCherry-U6-shBLTP2#1 or AAV-PHP.eB-EF1a-DIO-mCherry-U6-shBLTP2#2.

For projection-specific genetic manipulation experiments (Fig. 5b–d, l–q, and Fig. 6), spCas9 mice received bilateral mPFC injections of a total volume of 800 nL containing AAV-PHP.eB-CaMKII-Cre together with either AAV-PHP.eB-U6-sgRNA(Scramble)_hSyn-mCherry-KASH-bGH or AAV-PHP.eB-U6-sgRNA(BLTP2)_hSyn-mCherry-KASH-bGH. In additional experiments, C57BL/6J mice received bilateral mPFC injections of a total volume of 800 nL containing pAAV-pMecp2-SpCas9-spA together with either AAV-PHP.eB-U6-sgRNA(Scramble)_hSyn-mCherry-KASH-bGH or AAV-PHP.eB-U6-sgRNA(BLTP2)_hSyn-mCherry-KASH-bGH.

For electrophysiological experiments (Fig. 5e**–**k), C57BL/6J mice received bilateral mPFC injections of a total volume of 800 nL containing AAV-PHP.eB-CaMKII-Cre, pAAV-EF1a-DIO-hChR2(H134R)-P2A-EGFP and either AAV-PHP.eB-EF1a-DIO-mCherry-U6-shCont or AAV-PHP.eB-EF1a-DIO-mCherry-U6-shBLTP2#1. All mPFC injections were made at the following stereotaxic coordinates relative to bregma: anteroposterior (AP), +2.58 mm; mediolateral (ML), ±0.5 mm; dorsoventral (DV), −0.9 mm. For BioID and dual-eGRASP experiments involving the BLA, injections were made at the following coordinates relative to bregma: AP, −1.4 mm; ML, ±3.35 mm; DV, −4.4 mm.

Mice were group-housed (2–5 per cage) under a 12-h light/dark cycle with *ad libitum* access to food and water. Animals were excluded from analysis if *post hoc* histology revealed off-target injections, unilateral expression, or insufficient expression in the target region; exclusion criteria were pre-specified. Final sample sizes are reported in the figure legends and Supplementary Table 3.

### Immunocytochemistry and imaging analysis

Cultured neurons were transduced with small-scale AAVs at DIV 0 as previously described ^39^. Neurons were fixed at DIV 10 in 4% PFA for 20 min at room temperature. They were permeabilized with 0.1% Triton X-100 and 5% normal goat serum (NGS) for 30 min at room temperature. Samples were then incubated overnight at 4°C with primary antibodies followed by Alexa Fluor 488-, Alexa Fluor 555- or Alexa 647-conjugated secondary antibodies diluted in PBS containing 0.01% Triton X-100 and 5% NGS for 2 h at room temperature. The neuron and HEK293T cell mixed-culture assay was performed as previously described ^39^. Briefly, HEK293T cells were transfected with plasmid DNA (NGL-1-myc/pEGFP-N1) using Lipofectamine 2000 (Invitrogen) according to the manufacturer’s instructions. After 20 h, the cells were seeded on cultured neurons at DIV14. Fluorescence images were acquired on a Zeiss Imager microscope equipped with an Apotome 3 module, Zeiss 900 and Zeiss 980 confocal microscopes using ZEN software or a stimulated emission depletion (STED) super-resolution microscope (TCS SP8 STED, Leica Microsystems) using Leica Application Suite software. The individual acquiring the images was always blinded to the experiment. Images were quantified and post-processed using FIJI.

### Artificial synapse formation assay

The neuron–HEK293T mixed-culture assay for artificial synapse formation was performed as previously described with minor modifications. HEK293T cells were transfected with NGL-1-myc/pEGFP-N1 (Neuroligin-1 fused to EGFP) using Lipofectamine 2000 (Invitrogen) according to the manufacturer’s instructions. At 18–24 h after transfection, HEK293T cells were detached with 0.05% trypsin–EDTA (Gibco), resuspended in neuron culture medium and seeded onto cortical neuron cultures at DIV14 at a density of approximately 1–2 × 10^4^ cells per 12-mm coverslip. Co-cultures were maintained for 20–24 h at 37 °C in 5% CO₂ and then fixed in 4% PFA in PBS for 20 min at room temperature. For immunostaining, cells were permeabilized with 0.1% Triton X-100 in PBS containing 5% normal goat serum (NGS) for 30 min and incubated overnight at 4 °C with primary antibodies diluted in PBS containing 0.01% Triton X-100 and 5% NGS. The following primary antibodies were used: chicken anti-GFP to visualize EGFP-Neuroligin-1 and rabbit anti-Neurexin-1α. After washing, Alexa Fluor 488- or 555-conjugated secondary antibodies were applied for 1 h at room temperature. Coverslips were mounted in ProLong Diamond Antifade Mountant (ThermoFisher) and stored at 4 °C until imaging. All image acquisition and quantification were performed by an experimenter blinded to the experimental condition.

### Synaptic staining and imaging analysis

Immunohistochemistry was performed as previously described ^39,75^. Briefly, brains were fixed in 4% PFA, and coronally or sagittally sectioned with a micro slicer (D.S.K.) at a thickness of 50 μm or 100 μm. The sections were incubated with primary antibodies diluted in PBS containing 0.1% Triton X-100 and 5% NGS at 4°C overnight, followed by incubation with Alexa Fluor 488-, Alexa Fluor 555- or Alexa Fluor 647-conjugated secondary antibodies diluted in PBS with 0.1% Triton X-100 and 5% NGS for 2 h at room temperature. The nuclei were visualized by staining with DAPI. Colocalization analysis for BLTP2 localization and synaptic number quantification was performed as previously described ^39^. Briefly, P40 control and experimental tissue sections were immunostained with an antibody against BLTP2 together with antibodies against presynaptic marker VGLUT1 and the postsynaptic marker Homer1, which together label excitatory synapses. For synaptic analysis, 5-μm-thick Z-stacks consisting of 15 optical sections were acquired at high magnification using a Zeiss 980 confocal microscope (63× objective plus 2.5× optical zoom, ZEN software). Each Z-stack was then converted into maximum projection images (MPIs) by condensing three consecutive optical sections using ImageJ. The colocalization of VGLUT1/Homer1 signals was determined using the ImageJ Image Calculator, and the number of colocalized synaptic puncta was quantified with the ImageJ Analyze Particles tool. For each image, synaptic puncta were quantified within regions of interest (ROIs) of 100 μm^2^, carefully selected from areas lacking neuronal cell bodies to avoid background interference. All results of the statistical analysis are shown in Supplementary Table 3. All image analyses were conducted by an investigator blinded to the experimental conditions.

For the analysis of dual-eGRASP images, confocal image processing and 3D reconstruction of mPFC–BLA synapses were conducted using ZEN software. To prevent any bias in the analysis, all image samples were blinded before the evaluation. Cyan (CFP) and yellow (YFP) eGRASP signals were automatically detected using both ZEN and FIJI software. Yellow or yellow + cyan eGRASP signals were classified as activated synapses originating from the mPFC–BLA neural pathway, while cyan-only eGRASP signals were classified as synapses originating from the non-activated mPFC–BLA neural pathway. The identity of the postsynaptic neurons was determined based on the fluorescent proteins expressed on the cell membrane. Spine density data were exported to the experimenter in a blinded manner. The raw data for each dendrite were then paired with the corresponding GRASP information (cyan, yellow, or no GRASP). Dendrites lacking cyan eGRASP signals were excluded to ensure more precise analysis. Spine density was quantified within defined ROIs, typically 100 μm^2^ in size, carefully selected from areas devoid of neuronal cell bodies to minimize background interference. For each image, the number of synaptic puncta within each ROI was counted, and the density of spine synapses was calculated. All results of the statistical analysis are shown in Supplementary Table 3.

### Immunoprecipitation and immunoblotting

Co-immunoprecipitation experiments were performed using a HEPES-based buffer as described previously. Tissue was resuspended in HEPES buffer (20 mM HEPES, pH 7.5, 150 mM NaCl, 1 mM EDTA, 1 mM DTT, 0.2% NP-40) supplemented with protease and phosphatase inhibitor cocktails (cOmplete Mini, EDTA-free; PhosSTOP, Roche) and incubated for 1 h at 4 °C with gentle rotation. Lysates were sonicated on ice (3 × 10 s) and clarified by centrifugation at 20,400 × g for 10 min at 4 °C. Supernatants were transferred to low protein-binding tubes and precleared with Sepharose beads for 30 min at 4 °C with rotation. After centrifugation at 1,500 × g for 1 min, the supernatant was incubated overnight at 4 °C with rabbit anti-Neurexin 1 antibody. The following day, protein G–Sepharose beads were added, and samples were rotated for 1 h at 4 °C. Beads were collected by centrifugation at 1,500 × g for 1 min, and the supernatant was removed. Beads were washed five times with ice-cold wash buffer of the same composition as the lysis buffer. After the final wash, residual buffer was carefully removed, and bound proteins were eluted in Laemmli SDS sample buffer by heating at 95 °C for 5 min before SDS–PAGE and immunoblotting.

### Synaptosome fractionation

Synaptosome-enriched fractions were prepared from microdissected basolateral amygdala (BLA) using a sucrose step-gradient protocol. All procedures were carried out on ice or at 4 °C. BLA tissue was rapidly dissected in ice-cold PBS and transferred to 1.0 mL of homogenization buffer (0.32 M sucrose, 10 mM HEPES–NaOH, pH 7.4, 1 mM EDTA, protease and phosphatase inhibitors). Tissue was homogenized with a Dounce A glass homogenizer (20 strokes), and a 50-μL aliquot was saved as crude homogenate. The homogenate was centrifuged at 1,000 × g for 10 min at 4 °C, and the supernatant was collected into a new tube. The pellet (P1; containing nuclei and tissue debris) was resuspended in 1.0 mL homogenization buffer, homogenized again (20 strokes) and centrifuged at 1,000 × g for 10 min; the two supernatants were combined to yield the S1 fraction, which contains membranes and synaptosomes. S1 was centrifuged at 30,000 × g in a TLA55 rotor for 30 min at 4 °C to obtain the P2 pellet. The P2 pellet was gently resuspended in 500 μL of 0.32 M sucrose, 1 mM NaHCO₃ (pH 8.3) and homogenized with a Dounce B pestle (10 strokes); a 100-μL aliquot was retained as the total P2 fraction. For sucrose density-gradient separation, 400 μL of the P2 suspension was layered onto a discontinuous gradient consisting of 600 μL each of 0.85 M, 1.0 M and 1.2 M sucrose in 1 mM NaHCO₃ (TLS55 tubes) and centrifuged at 120,000 × g for 2 h at 4 °C. Three fractions were collected: the interface between 0.32 M and 1.0 M sucrose (myelin-enriched), the interface between 1.0 M and 1.2 M sucrose (synaptosome fraction) and the pellet (mitochondria-enriched). The synaptosome fraction (1.0–1.2 M interface) was diluted threefold with 30 mM Tris–HCl, pH 8.0, and centrifuged at 136,000 × g in a TLA55 rotor for 30 min at 4 °C. The resulting pellet was resuspended in 0.4 mL of 20 mM Tris–HCl, pH 8.0, and used immediately for downstream biochemical analyses or stored at −80 °C until use.

### SDS–PAGE and immunoblotting

SDS–PAGE and immunoblotting were performed as previously described ^39,75^, with minor modifications. Protein samples from whole lysates, immunoprecipitates or subcellular fractions were mixed with 2× or 4× Laemmli SDS sample buffer and heated at 95 °C for 5 min or 65 °C for 15 min. Proteins were separated by SDS–PAGE using Mini-PROTEAN electrophoresis systems (Bio-Rad) and transferred onto nitrocellulose membranes (Cytiva, Cat# 10600002) using a PowerPac HC power supply (Bio-Rad). Membranes were blocked for 1 h at room temperature in Tris-buffered saline containing 0.1% Tween-20 (TBS-T) and 5% skim milk, and then incubated overnight at 4 °C with primary antibodies diluted in TBS-T containing 5% skim milk. After washing in TBS-T, membranes were incubated for 1 h at room temperature with fluorophore-conjugated secondary antibodies diluted in TBS-T. Signals were detected using a ChemiDoc Touch MP imaging system (Bio-Rad) and quantified with Image Lab software (Bio-Rad) or FIJI. Band intensities were normalized to loading controls, and details of the quantification procedures are provided in the Statistical analysis section.

### In vivo BioID protein purification

*In vivo* BioID experiments were performed as previously described with some modifications ^39^. AAV-PHP.eB-CaMKII-ProX-ID virus was transduced into the mPFC of adult mouse brains (P45). Three weeks post-virus injection, biotin (24 mg/kg, s.c.) was administered and brains were collected 12 h later to enhance biotinylation efficiency. For each *in vivo* BioID experiment, a single mouse brain was used for each biotinylated protein purification, and each purification was performed independently at least four times. After each brain sample was lysed in a buffer containing 50 mM Tris-HCl (pH 7.5), 150 mM NaCl, 1 mM EDTA and a protease inhibitor mixture (cOmplete Mini EDTA-free, Roche), an equal volume of lysis buffer was added, containing 50 mM Tris-HCl (pH 7.5), 150 mM NaCl, 1 mM EDTA, 0.4% SDS, 2% Triton X-100, 2% deoxycholate, protease inhibitor mixture and phosphatase inhibitor mixture. The samples were then sonicated and centrifuged at 15,000 × *g* for 10 min. The supernatant was further ultracentrifuged at 100,000 × *g* for 30 min at 4°C. SDS was added to the cleared supernatant to a final concentration of 1%, and the sample was heated at 65°C for 15 min. After cooling on ice, the sample was incubated with Pierce High Capacity NeutrAvidin Agarose (ThermoFisher) at 4°C overnight. The beads were washed twice with 2% SDS, twice with 1% Triton X-100, 1% deoxycholate and 25 mM LiCl, twice with 1 M NaCl and five times with 50 mM ammonium bicarbonate. Biotinylated proteins were eluted in a buffer containing 125 mM Tris-HCl (pH 6.8), 4% SDS, 0.2% β-mercaptoethanol, 20% glycerol and 3 mM biotin at 65°C for 15 min.

### Purification of biotinylated proteins followed by mass spectrometry

Each sample was prepared by pooling BLA or PFC tissues from three mice and was lysed in 1.5 mL of guanidine-TCEP buffer (7 M guanidine-HCl, 100 mM HEPES-NaOH, pH7.5, 10 mM TCEP and 40 mM chloroacetamide). The lysates were heated, sonicated and centrifuged at 20,000 g for 15 min at 4°C. Supernatants were collected, and proteins (2 mg each) were purified by methanol–chloroform precipitation and solubilized in 150 µL of a buffer containing 8 M urea, 1% SDS, 50 mM Tris-HCl (pH7.5) and 150 mM NaCl. After sonication, the protein solutions were diluted eightfold with a buffer containing 50 mM Tris-HCl (pH7.5) and 150 mM NaCl. Biotinylated proteins were captured on a 10 µL slurry of NanoLink streptavidin magnetic beads (Vector) by incubation for 3 h at 4°C. The beads were washed three times with a buffer containing 1 M urea, 0.125% SDS, 50 mM Tris-HCl (pH7.5), and 150 mM NaCl, followed by three washes with 1 M urea and 50 mM ammonium bicarbonate. Proteins on the beads were digested with 400 ng of trypsin/Lys-C mix (Promega) overnight at 37°C. The digested peptides were acidified and desalted using GL-Tip SDB (GL Sciences). The eluates were evaporated in a SpeedVac concentrator and reconstituted in 0.1% trifluoroacetic acid and 3% acetonitrile (ACN).

LC-MS/MS analysis of the resulting peptides was performed using an EASY-nLC 1200 UHPLC coupled to an Orbitrap Fusion mass spectrometer via a nanoelectrospray ion source (Thermo Fisher Scientific). Peptides were separated on a 75 µm inner diameter × 150 mm C18 reversed-phase column (Nikkyo Technos) with a linear gradient of 4–32% acetonitrile for 0–100 min followed by an increase to 80% ACN for 10 min and a final hold at 80% ACN for 10 min. The mass spectrometer was operated in data-dependent acquisition mode with a maximum duty cycle of 3 s. MS1 spectra were measured with a resolution of 120,000, an AGC target of 4e5 and a mass range of 375 to 1,500 *m/z*. HCD MS/MS spectra were acquired in the linear ion trap with an AGC target of 1e4, an isolation window of 1.6 *m/z*, a maximum injection time of 35 ms, and a normalized collision energy of 30. Dynamic exclusion was set to 20 s. Raw data were analyzed directly against the SwissProt database restricted to *Mus musculus* using Proteome Discoverer version 2.5 (Thermo Fisher Scientific) for protein identification and label-free precursor ion quantification. Search parameters were as follows: (a) trypsin as an enzyme with up to two missed cleavages; (b) precursor mass tolerance of 10 ppm; (c) fragment mass tolerance of 0.6 Da; (d) cysteine carbamidomethylation as a fixed modification; and (e) protein N-terminal acetylation and methionine oxidation as variable modifications. Peptides were filtered with a false discovery rate of 1% using the Percolator node.

### Purification of biotinylated peptides followed by mass spectrometry

Brain regions (mPFC, BLA, NAc, Thal, HT and CTX) were lysed in 0.5 mL of guanidine-TCEP buffer as described above, each with four biological replicates. Proteins (2 mg each) were purified by methanol–chloroform precipitation and solubilized in 195 µL of PTS buffer (12 mM SDC, 12 mM SLS, 100 mM Tris-HCl, pH8.0). After sonication and heating, the protein solutions were diluted fivefold with 100 mM Tris-HCl (pH8.0) and digested with trypsin (MS grade, Thermo Fisher Scientific) overnight at 37°C. The resulting peptide solutions were diluted twofold with TBS (50 mM Tris-HCl, pH 7.5, 150 mM NaCl). Biotinylated peptides were captured on a 15 µL slurry of MagCapture HP Tamavidin 2-REV magnetic beads ^76,77^ (FUJIFILM Wako) by incubation for 3 h at 4°C. After washing with TBS five times, the biotinylated peptides were eluted with 100 µL of 1 mM biotin in TBS for 15 min at 37°C twice. The combined eluates were desalted, evaporated and reconstituted in 0.1% trifluoroacetic acid and 3% ACN.

LC-MS/MS analysis was performed on the same system as above. The peptides were separated with a linear 4–32% ACN gradient for 0–60 min, followed by an increase to 80% ACN for 10 min and a final hold at 80% ACN for 10 min. The MS1 spectra were acquired under the same settings as above. HCD MS/MS spectra were acquired in a linear ion trap with an AGC target of 1e4, an isolation window of 1.6 *m/z*, a maximum injection time of 200 ms and a normalized collision energy of 30. Dynamic exclusion was set to 10 s. Database searching was as above, except that lysine biotinylation was additionally set as a variable modification. Label-free quantification was performed based on the intensities of the precursor ions using the precursor ions quantifier node.

### Proteome data analysis

The proteomic dataset was further processed using Perseus (version 2.0.1.1). Proteins detected in fewer than three samples were removed. Normalized abundance values were log_2_-transformed, and differential protein biotinylation level between control and ProX-ID samples was evaluated using Student’s *t*-tests. The resulting *p*-values were corrected for multiple comparisons using the Benjamini–Hochberg false discovery rate (BH-FDR) method. To identify proteins significantly more enriched in the ProX-ID samples compared to controls, we applied a threshold of BH-FDR < 0.05 and a log_2_ fold change > 2. Proteins meeting these criteria were ranked by log_2_ fold change, and the top 200 were defined as a rank-based subset. For the mPFC–CTX projection, this subset comprised 176 proteins, reflecting the total number passing the significance thresholds. GO Cellular Component (GO CC) annotations of the identified proteins were assigned using the MGI (Mouse Genome Informatics) database, with classification based on the following terms: cytoplasm (GO:0005737), plasma membrane (GO:0005886), cytoskeleton (GO:0005856), mitochondrion (GO:0005739), Golgi apparatus (GO:0005794), endoplasmic reticulum (GO:0005783), endosome (GO:0005768), synaptic vesicle (GO:0008021) and presynaptic active zone (GO:0048786). UniProt IDs were first converted to MGI gene IDs using a UniProt-to-MGI conversion table downloaded from the UniProt website. Fisher’s exact test was performed for each GO CC term using the full list of UniProt IDs from the ID conversion table as the background reference set. Adjusted *p*-values were calculated using the BH-FDR method. Known presynaptic proteins were defined as those included in the presynaptic gene set retrieved as a list of MGI gene IDs using the getAllGenes4Compartment() function from the synaptome.db package ^43^. Neurological disease-associated genes were identified using the getGeneDiseaseByIDs() function for DO terms listed in Supplementary Table 2. Human Entrez Gene IDs obtained from synaptome.db were first converted to MGI gene IDs using the HGNC Comparison of Orthology Predictions (HCOP) database (https://www.genenames.org/tools/hcop/). The resulting MGI gene IDs were then converted to UniProt IDs using the UniProt-to-MGI conversion table described above, for subsequent comparison with our proteomic dataset. Proteins with no GO annotations, excluding those with low evidence codes (IC, IEA, NAS, ND, TAS), were classified as functionally uncharacterized proteins (GO). GO enrichment analysis was performed using Metascape (http://metascape.org). We specifically employed the Custom Analysis workflow to refine our analysis to select ontology catalogues including Biological Process and Molecular Function. The background was defined as the default species genome (*M. musculus*). The enrichment parameters were set to default thresholds: minimum 3 genes per term, *p* ≤ 0.01 and enrichment factor ≥ 1.5. Dot plots were generated using the output from Metascape and visualized in MATLAB. The plots display the –log₁₀(*p*-value) and the percentage of annotated proteins relative to the total input for the top 20 enriched terms. Protein–term networks were generated based on annotations from the KEGG BRITE database ^46–48^. For visualization, the KEGG BRITE terms in the following categories were used: BRITE Hierarchy (09180) and Environmental Information Processing (09130) in KEGG Orthology [BR:mmu00001] (https://www.kegg.jp/brite/mmu00001), Membrane Trafficking [BR:mmu04131] (https://www.kegg.jp/brite/mmu04131), Ion Channels [BR:mmu04040] (https://www.kegg.jp/brite/mmu04040), Transporters [BR:mmu02000] (https://www.kegg.jp/brite/mmu02000) and GTP-Binding Proteins [BR:mmu04031] (https://www.kegg.jp/brite/mmu04031). To improve visual clarity, some categories were merged and manually labelled. For proteins associated with multiple terms, a subset of associations was not displayed to maintain the interpretability of the network. Protein–protein interaction (PPI) networks were constructed using the Reactome FI plugin in Cytoscape ^78^. PPI edges for all human genes were initially retrieved, and the corresponding human gene symbols were converted to MGI IDs using the HCOP database. These MGI IDs were subsequently mapped to UniProt IDs using the UniProt–MGI ID conversion table, as described above.

### Mouse single-nucleus RNA sequencing

Single-nucleus RNA sequencing (snRNA-seq) data were obtained from the Allen Brain Cell Atlas (https://alleninstitute.github.io/abc_atlas_access/intro.html) ^79^. The log-normalized AnnData object containing mPFC layer 2/3 neurons was downloaded via the *abc_atlas_access* Python library (v0.1.2). For comparison, the cluster annotated as *007_L2/3_IT_CTX_Glut* in the original publication was extracted, and mean expression levels and expression proportions of genes identified from the mPFC–BLA proteome were calculated using the *Scanpy* Python library ^80^. All analyses were performed in Python (3.10.11).

### Marmoset spatial transcriptomics

Marmoset 10x Visium data were obtained from a previous study ^81^. The mPFC was manually delineated based on H&E staining patterns. Analyses were conducted using the *Seurat* R package (v5.3.0) ^82^. The data were processed with the following *Seurat* functions: *SCTransform* for normalization, *RunPCA* for dimensionality reduction, *FindNeighbors* for computation of the nearest neighbors and *FindClusters* for cluster identification. Cortical layer 2 and layer 3 clusters were then defined based on characteristic cortical layer markers in combination with their spatial localization. After batch correction using *Harmony* R package ^83^, the mean expression levels of genes identified from the mPFC–BLA proteome were calculated using the *Seurat* R package. All analyses were performed in R (4.4.1).

### Behavioural analysis

BLTP2-cKO mice and control mice were subjected to a behavioural battery in the following order: open field test, elevated plus maze (EPM) test, three chamber social interaction test, and cued and contextual fear conditioning test. The open field, EPM and three-chamber tests were conducted in a sound-attenuated room, where mice were also habituated for 1 h prior to each test. The fear conditioning test was conducted in a conditioning chamber enclosed within a sound-attenuating box, and, prior to testing, mice were habituated for 1 h in a separate sound-attenuated room distinct from the testing chamber. All apparatuses were cleaned between animal trials using slightly acidic electrolyzed water to eliminate olfactory cues, except for the grid floor of the fear conditioning chamber, which was cleaned with 70% ethanol. Behavioural testing was performed between 9:00 a.m. and 6:00 p.m., with the order of testing counterbalanced across groups. Each behavioural test was conducted on a separate day. Mouse behaviours were recorded using a video camera and analyzed by the SMART 3.0 video tracking system (Panlab Harvard Apparatus, Spain).

*Open field test.* The apparatus consisted of a white square cage (50 × 50 × 30 cm; O’Hara) with the floor divided into nine equal squares and illuminated at 40 lux. Each mouse was placed in the bottom-left corner of the arena and allowed to explore freely for 10 min. Behavioural activity was recorded, and the following parameters were quantified: total distance traveled (cm) and time spent in the center area (s).

*EPM test.* The apparatus consisted of two arms (25 × 5 cm) with 3-mm-high ledges along the sides and distal end (open arms) and two closed arms (25 × 5 cm) with transparent walls (15 cm high) enclosing the sides and distal ends (Shin Factory). Arms of the same type were positioned opposite each other and connected by a central square platform (5 × 5 cm), with all surfaces made of grey plastic. The maze was elevated 50 cm above the floor and illuminated at 100 lux at the center. Each mouse was placed in the central square facing a closed arm and allowed to explore for 10 min. Behavioural activity was recorded, and the following parameters were quantified: total distance traveled (cm) and time spent in the open arms (s).

*Three-chamber social interaction test.* The apparatus comprised a three-chamber rectangular structure with grey walls and floors and transparent dividing walls (Shin Factory). Each chamber measured 20 × 40 × 22.5 cm and was connected by walls containing small rectangular openings (5.5 × 8.5 cm) to allow free movement between chambers. On the day prior to testing, mice were habituated for 10 min in the central chamber, followed by 10 min of free exploration across all three chambers. On the test day, mice were again placed in the central chamber for 5 min and then allowed to explore all chambers freely for another 5 min. Following this exploration, a novel, age- and sex-matched C57BL/6J mouse (referred to as the stranger mouse) was placed inside a wire cage in one of the side chambers, while an identical but empty wire cage was placed in the opposite chamber. The test mouse was allowed to explore freely for 10 min. Behavioural activity was recorded, and the following parameters were quantified: total distance traveled (cm), time spent in the zone surrounding the cage containing the stranger mouse (social interaction time, s), and time spent in the zone surrounding the empty cage (empty cage time, s). A social interaction score was calculated as: (social interaction time − empty cage time)/(social interaction time + empty cage time).

*Fear conditioning test.* Each mouse was placed in a chamber (19 × 20 × 25 cm; context A) with a black wall, enclosed in a sound-attenuating box and illuminated at 200 lux. After 4 min of free exploration, the mice were exposed to a 30-s auditory tone (8 kHz, 70 dB; conditioned stimulus, CS) that coterminated with a 2-s foot shock (0.3 mA; unconditioned stimulus, US). This CS–US pairing was repeated three times with 130-s interstimulus intervals. Mice were removed from the chamber 130 s after the final pairing. Freezing behaviour during this final 130-s period was measured to assess fear acquisition. Cued and contextual tests were performed 1 day and 2 days after the conditioning, respectively. For the cued test, mice were placed in a novel environment (context B; 18 × 18 × 15 cm, white chamber with white walls) for a total of 8 min: a 3-min baseline period without the CS, a 3-min CS presentation (8 kHz, 70 dB) and a 2-min post-CS period. Freezing behaviour during the CS presentation period was measured to assess cued memory retrieval. For the contextual test, mice were returned to the original conditioning chamber (context A) and observed for 12 min without any stimuli. Freezing behaviour during the 12-min period was measured to assess contextual memory retrieval. The conditioning apparatus was used with Packwin software (Panlab Harvard Apparatus) on a PC. Freezing, defined as the complete absence of body movement except for respiration lasting at least 2 s, was automatically analysed using the SMART system (Panlab Harvard Apparatus).

### Electrophysiological analysis

Coronal brain slices containing the BLA were prepared using the same method previously described for the central amygdala ^84^. In brief, the mice were anaesthetized with 2% isoflurane and transcardially perfused with ice-cold cutting artificial cerebrospinal fluid (aCSF) (2.5 mM KCl, 0.5 mM CaCl_2_, 10 mM MgSO_4_, 1.25 mM NaH_2_PO_4_, 2 mM thiourea, 3 mM sodium pyruvate, 92 mM N-methyl-D-glucamine, 20 mM HEPES, 12 mM N-acetyl-L-cysteine, 25 mM D-glucose, 5 mM L-ascorbic acid and 30 mM NaHCO_3_ and pH 7.3–7.4 adjusted with HCl). Two to three minutes after perfusion, the brain was dissected out and cut into coronal slices containing the BLA. Coronal slices containing the BLA (300 μm thick) were made with a vibrating microtome (VT1200; Leica, Wetzlar, Germany), and then the slices were kept in the ice-cold aCSF. The slices were kept in a holding chamber with a constant flow of aCSF at 38°C for 10–15 min. They were then transferred to another holding chamber containing standard oxygenated aCSF solution (125 mM NaCl, 2.5 mM KCl, 2 mM CaCl_2_, 1 mM MgCl_2_, 1.25 mM NaH_2_PO_4_, 26 mM NaHCO_3_ and 20 mM glucose) at room temperature (22–25°C) for at least 30 min until electrophysiological recordings.

Slices were transferred to the recording chamber, and oxygenated aCSF solution (26–28°C) was continuously superfused at 4–7 mL/min. The patch pipettes were filled with an internal solution (125 mM K-gluconate, 10 mM KCl, 0.5 mM EGTA, 10 mM HEPES, 4 mM ATP-Mg, 0.3 mM NaGTP, 10 mM phosphocreatine, 5 mM QX-314, pH 7.28 adjusted with KOH). The pipette tip resistance was 4–9 MΩ. Synaptic currents were recorded using a computer-controlled amplifier (Axopatch 700B, Molecular Devices). The data were digitized with an analogue-to-digital converter (Digidata 1550, Molecular Devices), stored on a personal computer using a data acquisition program (pCLAMP 10.4 acquisition software, Molecular Devices), and analyzed using a software package (Clampfit version 10.7, Molecular Devices; MiniAnalysis, Synaptosoft). sEPSCs were recorded in the voltage-clamp mode at a holding potential −70 mV for 5 min. The frequency and amplitude of sEPSCs for 1 min were quantified using Minianalysis software (Synaptosoft). Light-evoked EPSCs (leEPSCs) were recorded in the voltage-clamp mode at a holding potential −70 mV. In the recording of leEPSCs, the slice was illuminated with a blue laser diode (COME2-LB473/532/100, Lucir: wavelength, 470 nm) and optical stimuli (intensity, 10 mW; duration, 5 ms) were applied to the recording area.

### Statistical analysis

Statistical analyses were performed exclusively on experimental data using GraphPad Prism v9 (GraphPad Software). Data are expressed as the mean ± SEM. We compared independent sample means using Student’s *t*-tests, one-way ANOVAs and two-way ANOVAs as appropriate. Statistically significant F-values detected in the ANOVAs were followed by alpha-adjusted *post-hoc* tests (Tukey’s HSD or Bonferroni’s test). *P* < 0.0001, *P* < 0.001, *P* < 0.01 and *P* < 0.05 were considered to indicate statistical significance. Sample size is indicated in the figure legend for each experiment. Sample sizes were determined based on previous experience for each experiment to yield high power to detect specific effects. No statistical methods were used to predetermine sample size. All results of the statistical analysis are shown in Supplementary Table 3.

### Implementation

Proteome annotation analysis was performed using MATLAB 2024a (MathWorks) and R 4.5.0. Visualization of protein networks was performed using Cytoscape 3.10.3 ^85^.

---

**Extended Data Figure 1.**
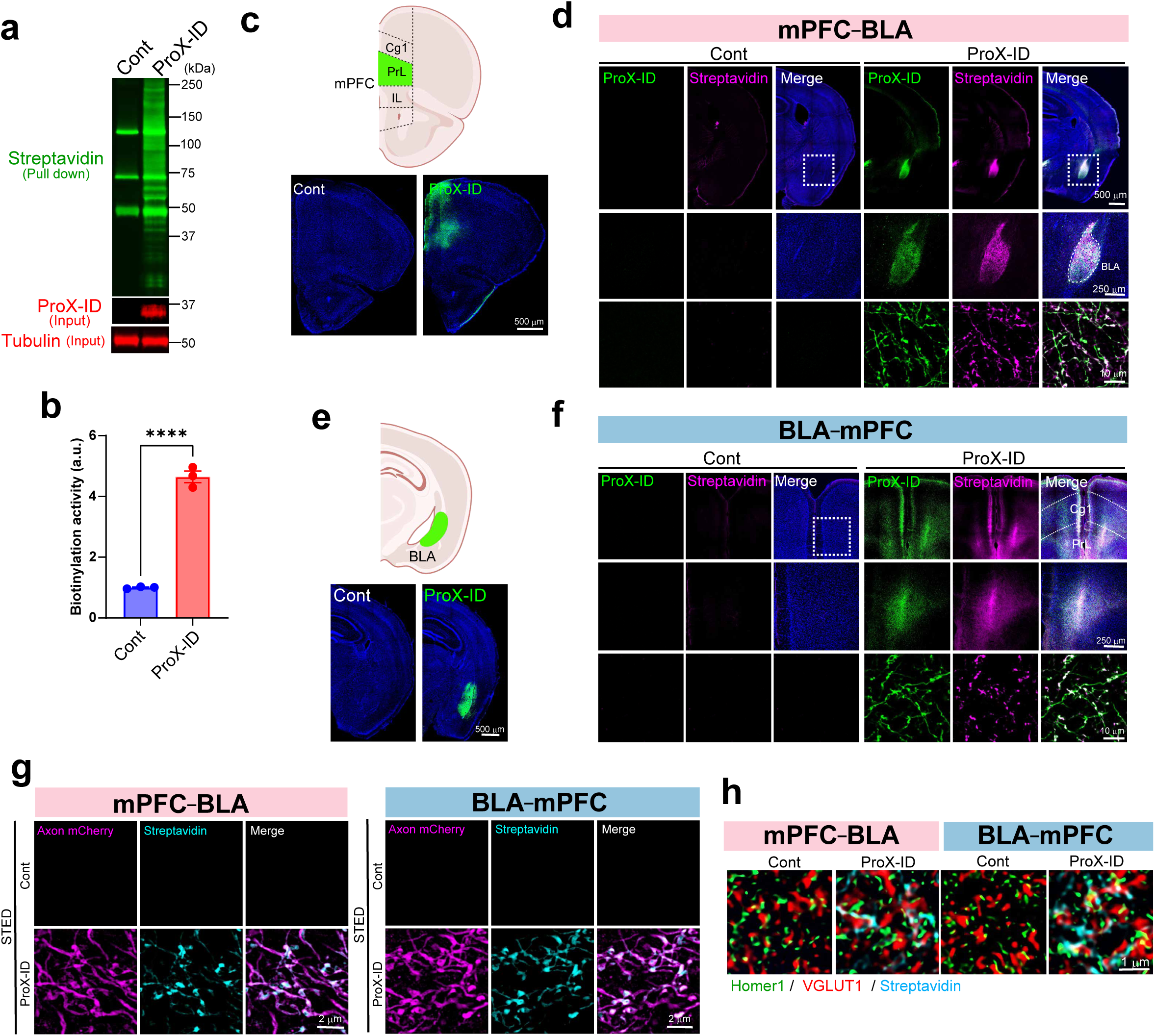
High-resolution and pathway-specific labelling of presynaptic terminals using ProX-ID. **a,** Representative images showing biotinylation activity in cultured neurons expressing ProX-ID at 14 days *in vitro* (DIV), confirming the increased biotinylation compared to control neurons. **b,** Quantification of streptavidin signals. Data are presented as mean ± s.e.m.; *n* = 4 per experimental group. Student’s *t*-test. \*\*\*\**P* < 0.0001. **c**–**d,** Immunohistochemistry of biotinylated proteins in the mPFC (**c**, injection site) and BLA (**d**, projecting site) illustrating labelling of the mPFC–BLA neural pathway. **e**–**f,** Immunohistochemistry of biotinylated proteins in the BLA (**e**, injection site) and mPFC (**f**, projecting site) illustrating labelling of the BLA–mPFC neural pathway. **g,** Super-resolution microscopy images of biotinylated proteins at axonal terminals of the mPFC–BLA (left) and BLA–mPFC (right) projections. **h,** Super-resolution microscopy images of biotinylated proteins with the excitatory synaptic markers Homer1 and VGLUT1.

**Extended Data Figure 2.**
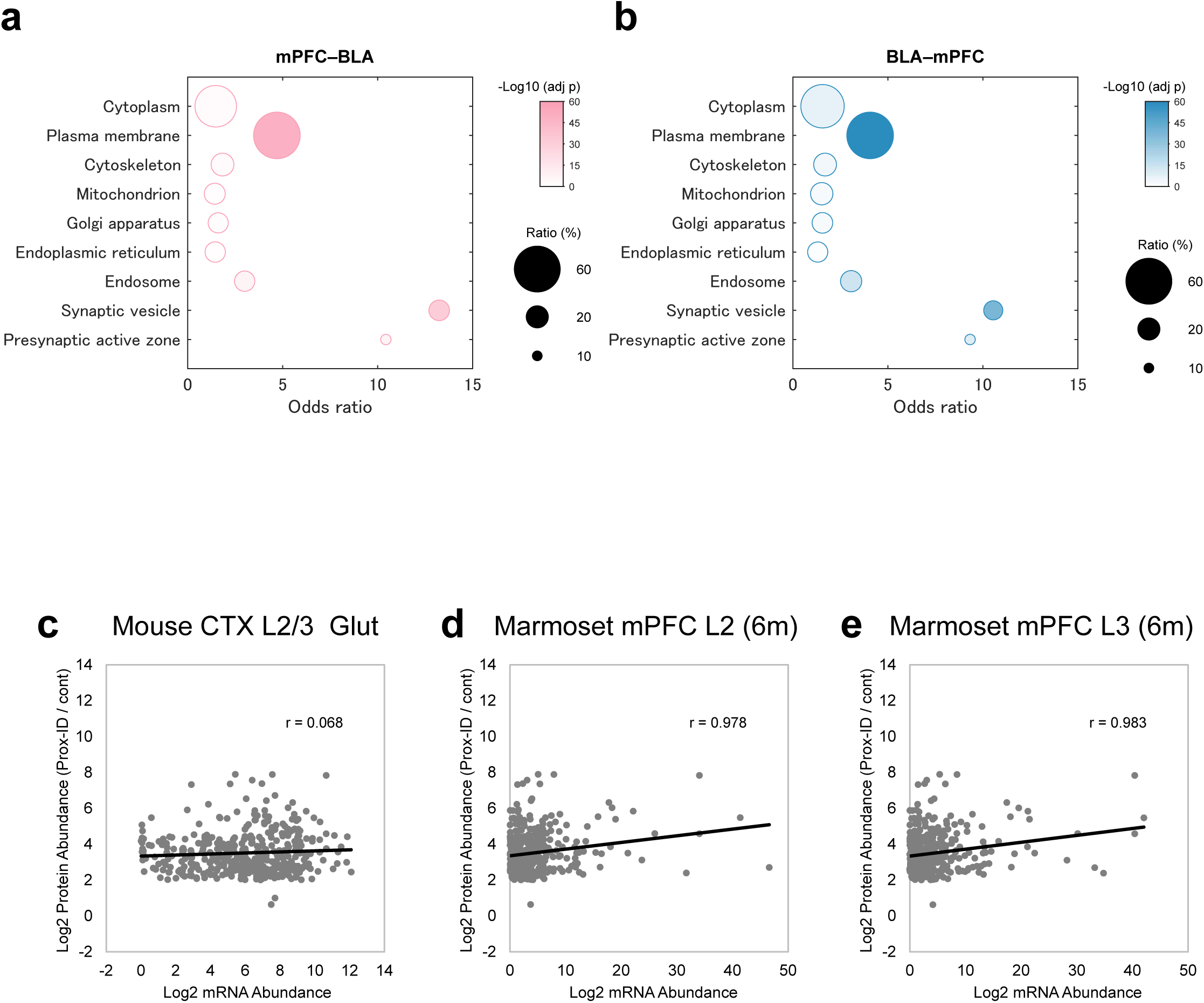
Presynaptic identity and transcriptomic concordance of mPFC–BLA projection-enriched proteins. **a**–**b,** Bubble plot summarizing enrichment analysis for the selected Gene Ontology (GO) Cellular Component terms (see Materials and Methods) based on Fisher’s exact test. **c**–**e,** Scatter plots showing gene expression levels in mouse (**c**) and marmoset **d**–**e,** mPFC neurons, plotted against protein biotinylation levels (ProX-ID/control) from mPFC–BLA samples. Gene expression levels in mouse layer 2/3 cortical neurons were obtained from mouse single-nucleus and spatial transcriptomic datasets in the Allen Brain Atlas. Gene expression levels in adult (six-month-old) marmoset mPFC layer 2 (**d**) and layer 3 (**e**) neurons were obtained from spatial transcriptomic analysis (see Materials and Methods).

**Extended Data Figure 3.**
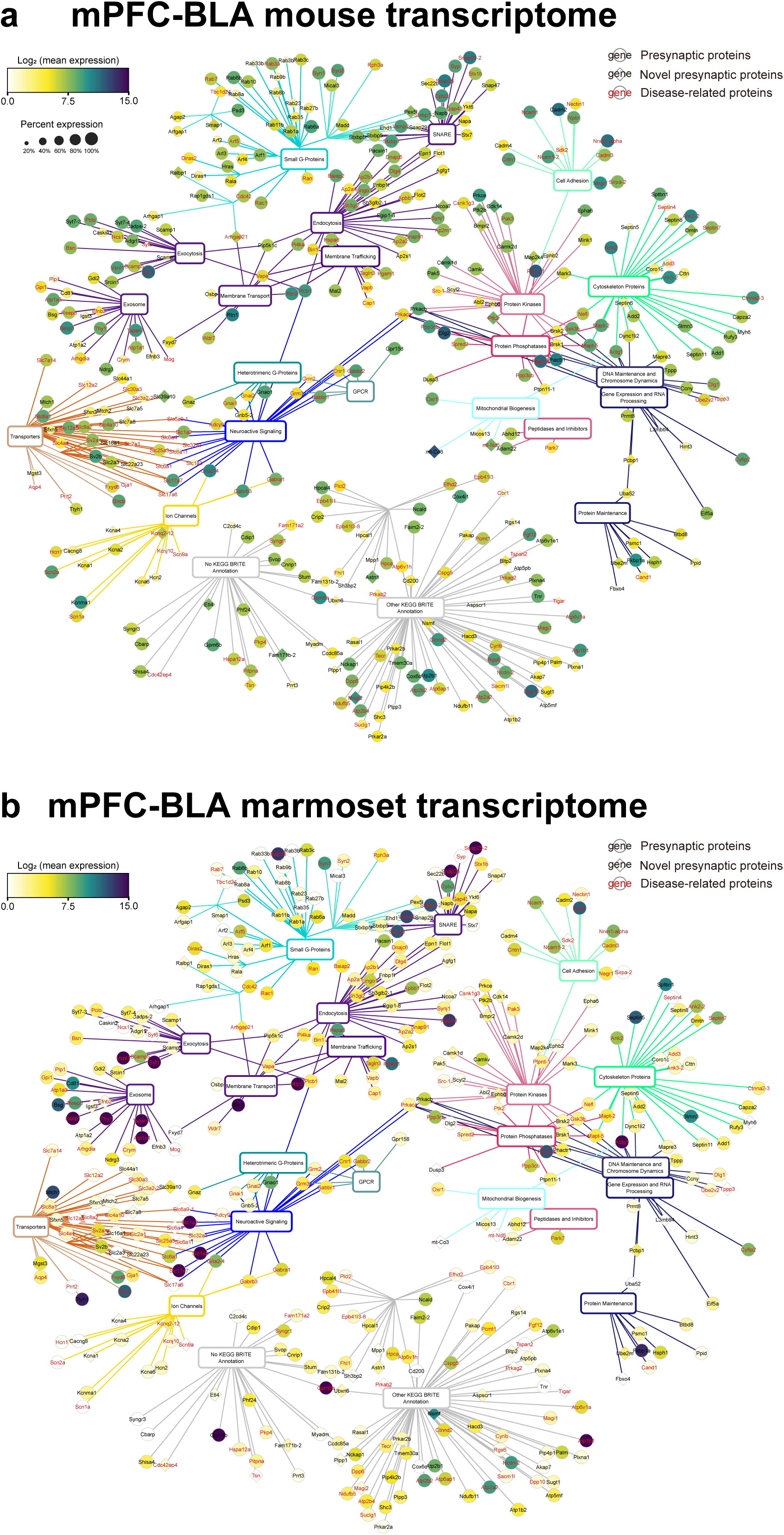
Cross-species transcriptomic mapping of projection-enriched synaptic proteins in mouse and marmoset cortex. Mouse (**a**) and marmoset (**b**) gene expression data for mPFC–BLA-enriched proteins were visualized in protein–term networks. The gene expression data shown in (**a**) and (**b**) correspond to the datasets presented in Extended Data Figures 2c and 2d, respectively.

**Extended Data Figure 4.**
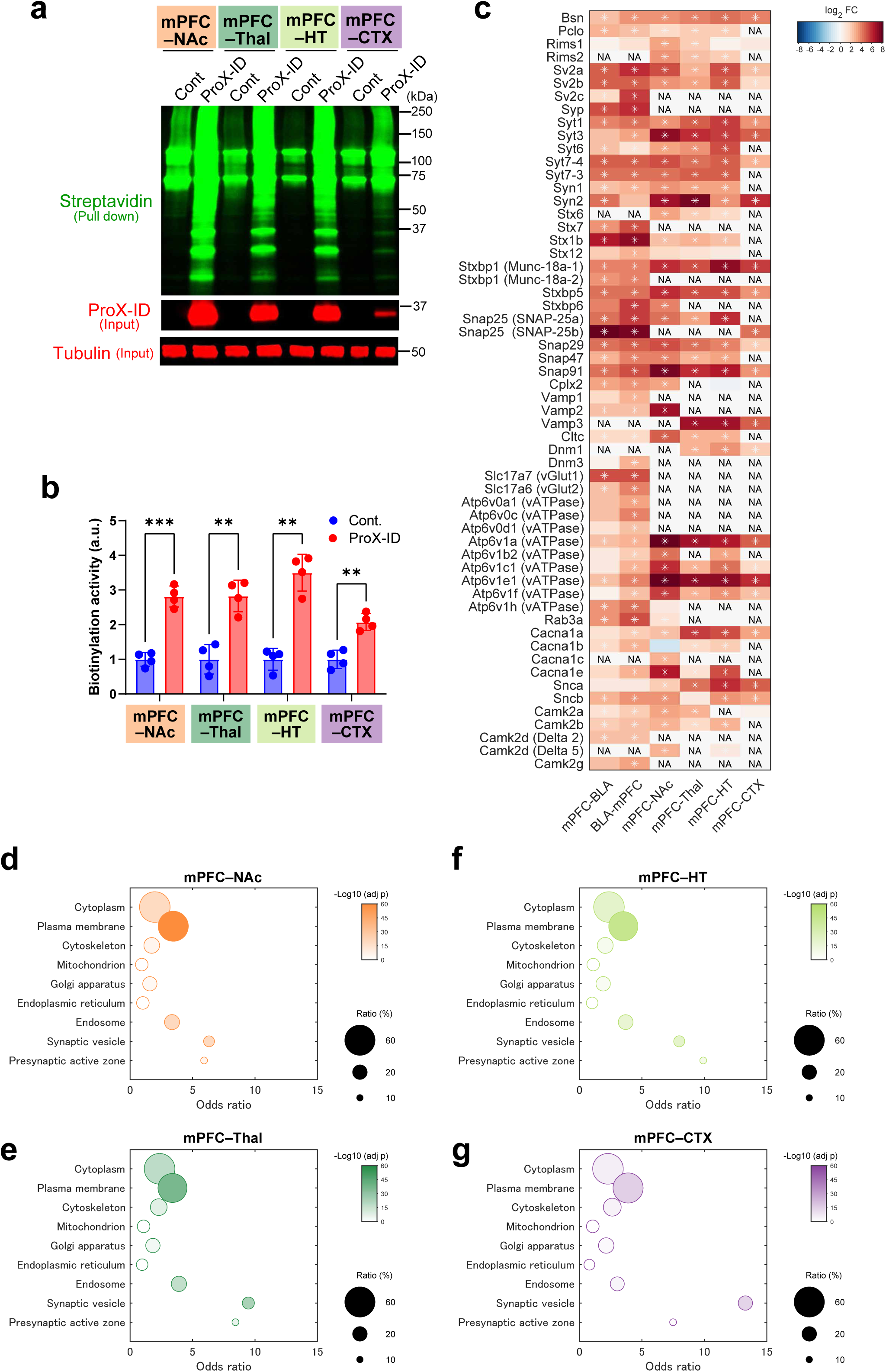
Quantitative profiling of presynaptic proteins across multiple mPFC output pathways. **a,** Immunoblot analysis of biotinylated proteins in the mPFC–NAc, mPFC–Thal, mPFC–HT and mPFC–CTX neural pathways. **b,** Quantification of streptavidin signals from NAc, Thal, HT and CTX lysates. Data are presented as mean ± s.e.m.; *n* = 4 mice per experimental group. Two-way ANOVA followed by Bonferroni *post hoc* test. \*\*\**P* < 0.001; \*\**P* < 0.01. Results for the selected pairwise comparisons are shown. **c,** Heatmap showing the normalized log_2_ biotinylation levels in ProX-ID samples relative to control samples for representative presynaptic proteins for mPFC–NAc, mPFC–Thal, mPFC–HT and mPFC–CTX together with mPFC–BLA, and BLA–mPFC. Asterisks indicate proteins significantly enriched in ProX-ID samples (FDR < 0.05, log_2_ fold change > 2, ProX-ID vs. control). NA indicates missing values. **d**–**g,** Bubble plots summarizing enrichment analysis for the selected Gene Ontology (GO) Cellular Component terms (see Materials and Methods) based on Fisher’s exact test.

**Extended Data Figure 5.**
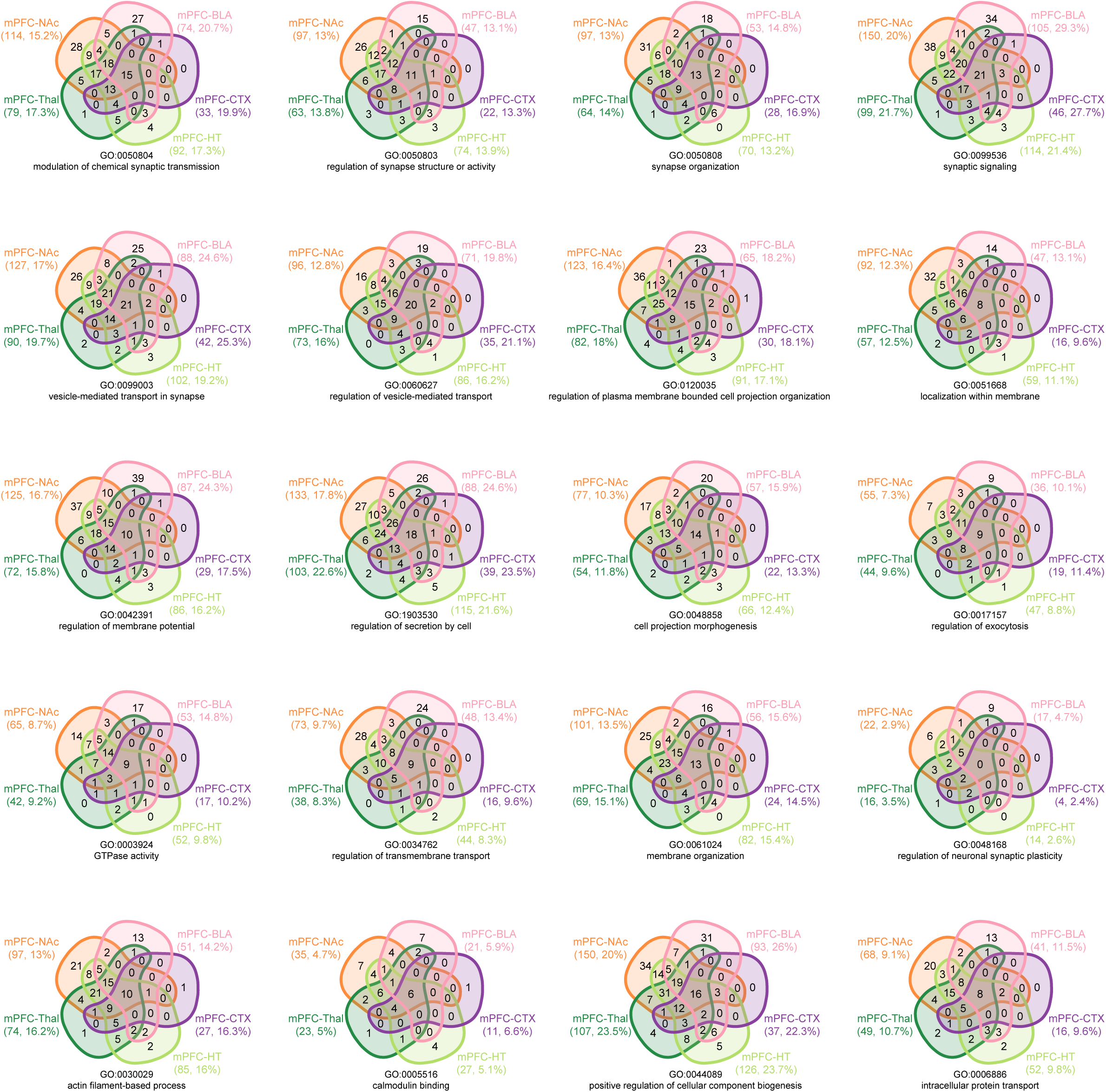
Divergence and overlap of synaptic functions across mPFC efferent pathways. Venn diagrams showing the overlap of neural pathway-enriched proteins among the mPFC–BLA, mPFC–NAc, mPFC–Thal, mPFC–HT and mPFC–CTX neural pathways. Numbers indicate the number of proteins in each intersection. The numbers in parentheses below each projection indicate the number of proteins annotated in the GO terms, and the percentages represent their ratio relative to the total proteins in each projection.

**Extended Data Figure 6.**
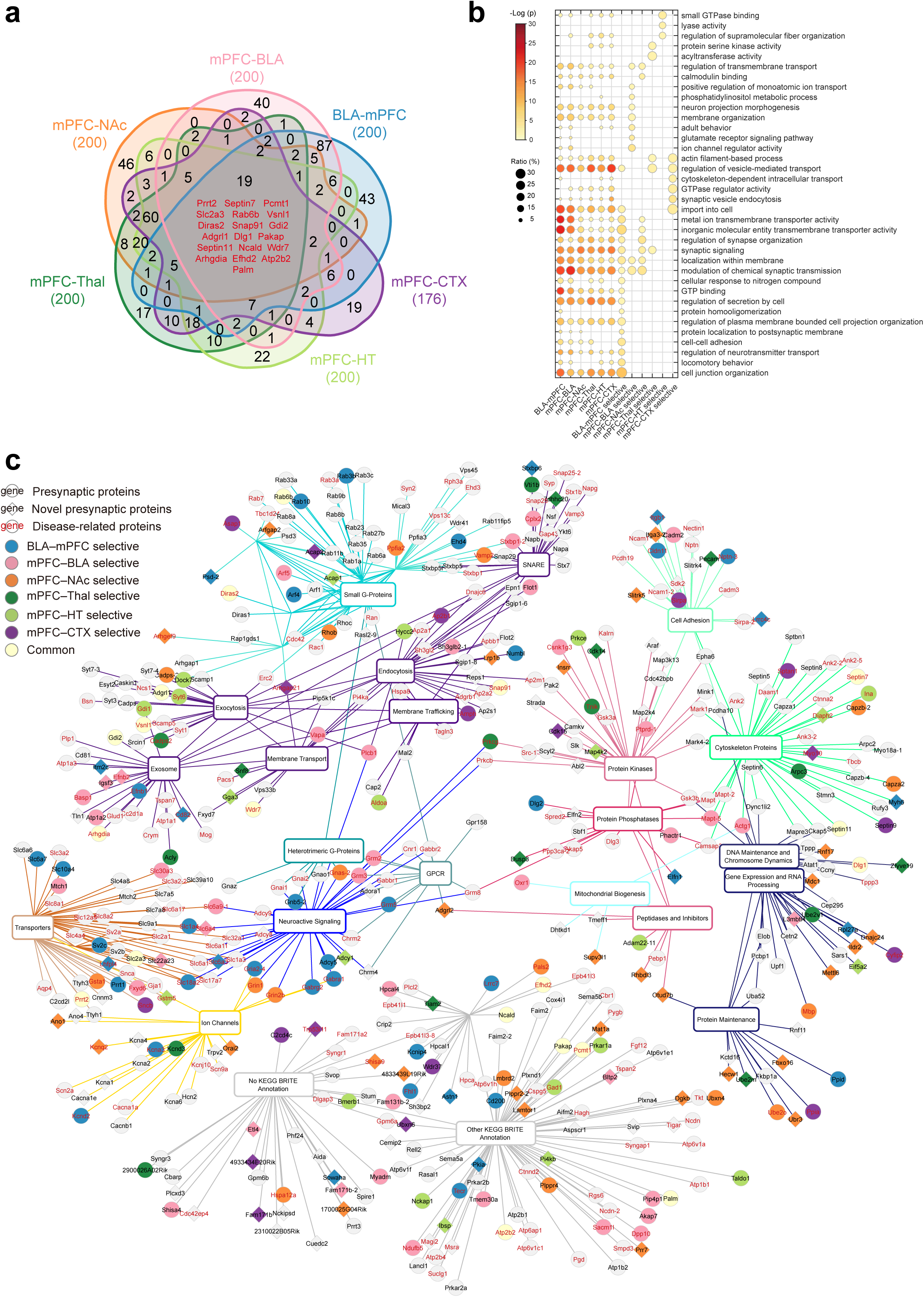
Comparison of the top 200 proteins ranked by log_2_ fold change further substantiates the molecular heterogeneity across the six mPFC-centric projection pathways. **a,** Venn diagram showing the overlap of significantly enriched proteins across mPFC–BLA, mPFC–NAc, mPFC–Thal, mPFC–HT, mPFC–CTX and BLA–mPFC neural pathways. **b,** Enriched GO terms for Biological Process and Molecular Function. **c,** KEGG BRITE-based protein–term network for projection enriched proteins (top 200 by log_2_ fold change). Nodes, proteins/terms; edges, associations based on BRITE functional categories.

**Extended Data Figure 7.**
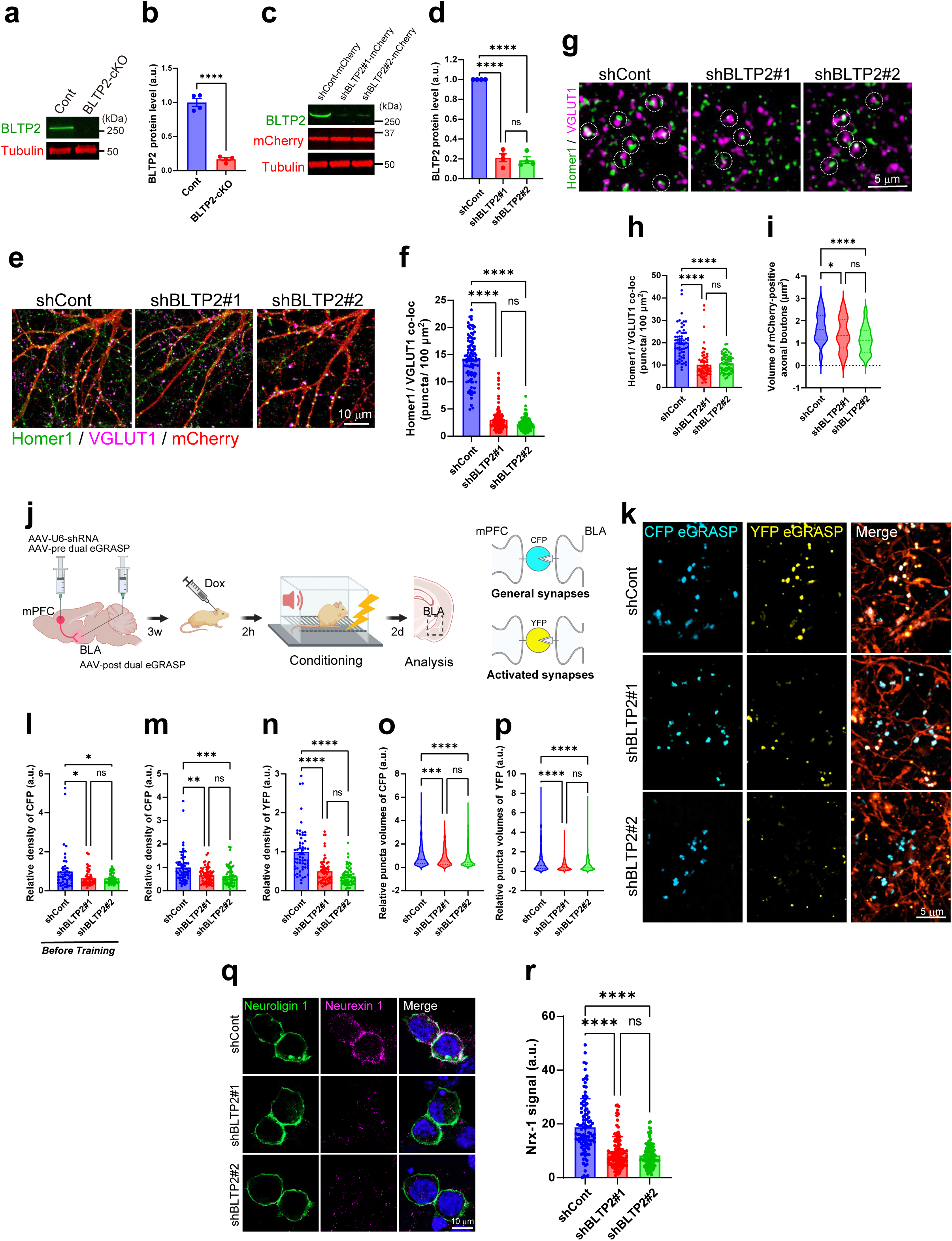
BLTP2 knockdown disrupts synaptic architecture and Neurexin 1 localization in the mPFC–BLA circuit. **a, c,** Immunoblot showing protein levels of BLTP2 in knockout (**a**) and knockdown (**c**) samples. **b**, **d,** Quantification of BLTP2 protein levels corresponding to (**a**) and (**c**), respectively. Data are presented as mean ± s.e.m.; *n* = 4 mice per experimental group. Student’s *t*-test for (**b**) and one-way ANOVA followed by Tukey’s *post hoc* test for (**d**). \*\*\*\**P* < 0.0001; ns, not significant. **e**–**f,** Quantification of the synaptic density of Homer1 and VGLUT1 double-positive excitatory synapses in cultured neurons after transfection with AAV-PHP.eB-U6-shCont or shBLTP2 (#1 or #2). Bar graph represents mean ± s.e.m.; shCont *n* = 96, shBLTP2#1 *n* = 108, shBLTP2#2 *n* = 120 cells. One-way ANOVA followed by Tukey’s *post hoc* test. \*\*\*\**P* < 0.0001; ns, not significant. **g,** Representative images of BLA sections from control and BLTP2-knockdown mice. Upper panels show immunostaining for Homer1 (green) and VGLUT1 (magenta). **h,** Quantification of the synaptic density of Homer1 and VGLUT1 double-positive excitatory synapses in the BLA. Data are presented as mean ± s.e.m.; *n* = 20 slices per group from each of four mice. One-way ANOVA followed by Tukey’s *post hoc* test. \*\*\*\**P* < 0.0001; ns, not significant. **i,** Quantification of presynaptic bouton volume in the BLA. Violin plots display the distribution of individual synaptic bouton volumes. The bold dashed line denotes the median, and the thin dashed lines indicate the first and third quartiles. *n* = 20 slices per group from each of four mice. One-way ANOVA followed by Tukey’s *post hoc* test. \*\*\*\**P* < 0.0001; \**P* < 0.05; ns, not significant. **j,** Schematic of the dual-eGRASP strategy combined with BLTP2-knockdown. **k,** Representative images of CFP- and YFP-eGRASP-labelled synapses in the BLA following fear conditioning in control and BLTP2 knockdown mice. **l–n,** Quantification of the synaptic density of CFP eGRASP-labelled and YFP eGRASP-labelled synapses. Data are presented as mean ± s.e.m.; *n* = 10 (**l**) or 20 (**m**, **n**) slices per group from each of four mice. One-way ANOVA followed by Tukey’s *post hoc* test. \*\*\*\**P* < 0.0001; \*\*\**P* < 0.001; \*\**P* < 0.01; \**P* < 0.05; ns, not significant. **o–p,** Quantification of the synaptic volume of CFP eGRASP-labelled (**o**) and YFP eGRASP-labelled (**p**) synapses after fear conditioning. Violin plots display the distribution of individual synaptic bouton volumes. The bold dashed line denotes the median, and the thin dashed lines indicate the first and third quartiles. shCont *n* = 511 (CFP) and 579 (YFP), shBLTP2#1 *n* = 465 (CFP) and 610 (YFP), shBLTP2#2 *n* = 534 (CFP) and 540 (YFP) synapses from each of four mice. \*\*\*\**P* < 0.0001; \*\*\**P* < 0.001; ns, not significant. **q,** Representative images showing subcellular localization of Neuroligin 1 (green) and Neurexin 1 (magenta) in neurons expressing shCont, shBLTP2#1 or shBLTP2#2. **r,** Quantification of Neurexin 1 signal intensity in Neuroligin 1-positive regions. Data are presented as mean ± s.e.m.; shCont *n* = 100, shBLTP2#1 *n* = 123, shBLTP2#2 *n* = 117 cells. One-way ANOVA followed by Tukey’s *post hoc* test. \*\*\*\**P* < 0.0001; ns, not significant.

**Extended Data Figure 8.**
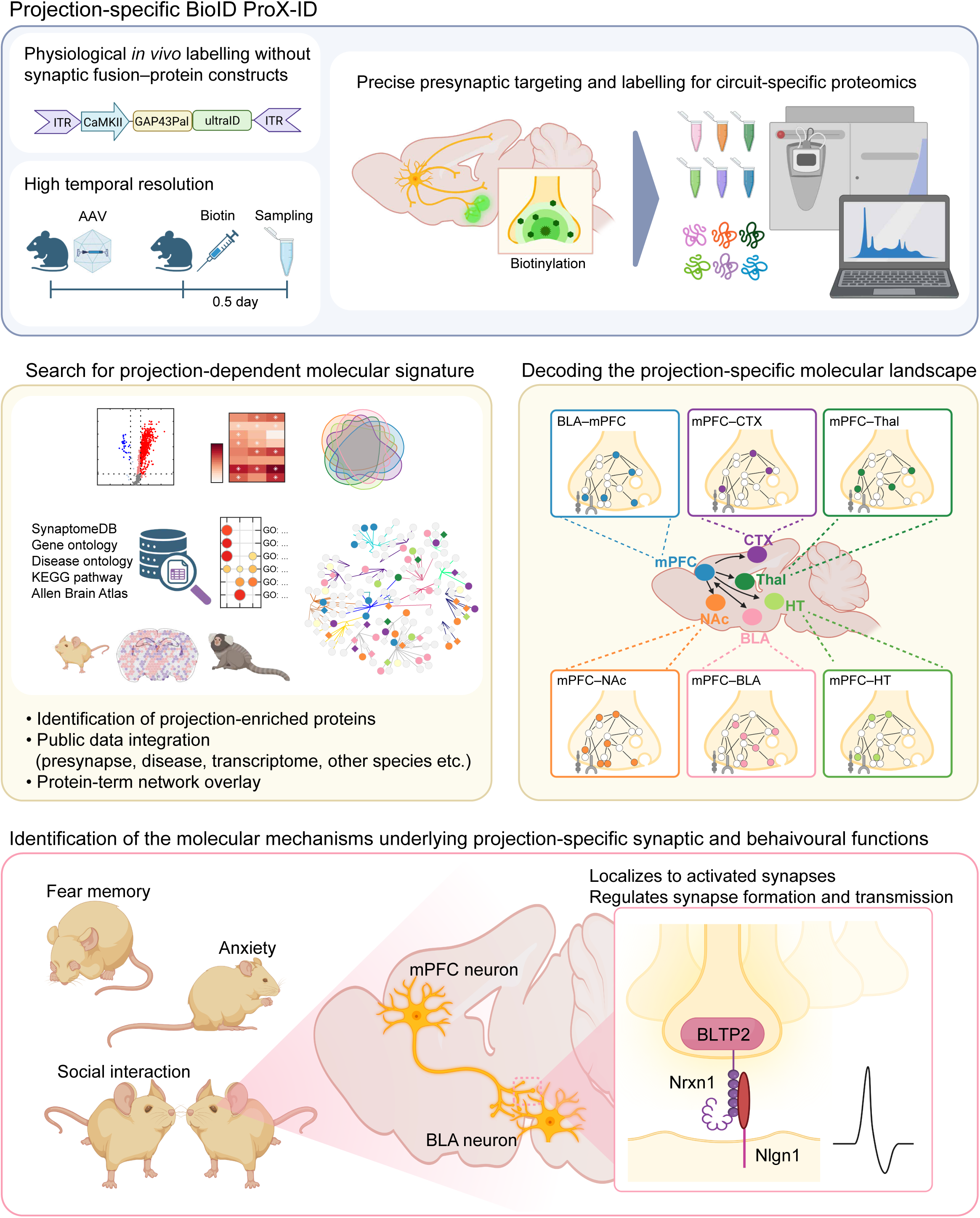
ProX ID maps projection specific presynaptic proteomes across mPFC centred pathways and links a BLTP2 Neurexin 1 axis to socioemotional behaviours Graphic summary of ProX ID, which enables physiological *in vivo* proximity labelling of presynaptic terminals with high temporal resolution, followed by streptavidin enrichment and quantitative mass spectrometry to derive pathway enriched presynaptic proteomes. Projection resolved signatures across six mPFC centred pathways (mPFC to BLA, NAc, Thal, HT and CTX, and BLA to mPFC) are integrated with public resources and cross species analyses to decode projection dependent molecular landscapes. The summary highlights BLTP2 enrichment in the mPFC to BLA pathway, its association with memory activated synapses, and a BLTP2 dependent presynaptic program involving Neurexin 1 that shapes synaptic organization and transmission, with consequences for fear memory, anxiety like behaviour and social interaction. Together, these findings reveal projection specific presynaptic molecular diversity and provide a mechanistic framework for circuit level vulnerabilities implicated in neuropsychiatric disorders.

**Supplemental Figure 1.**
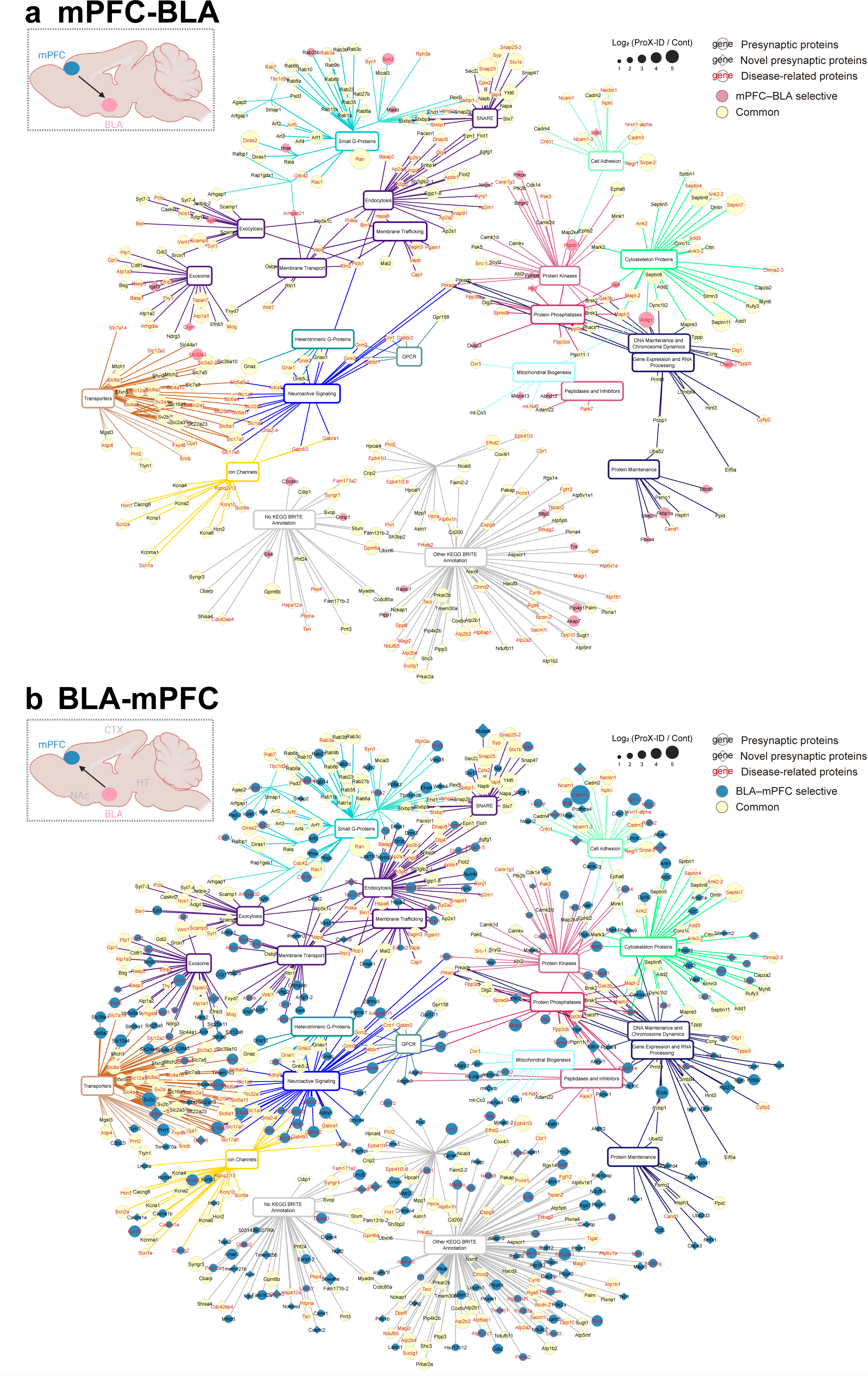
Protein–term network for neural pathway-enriched proteins. Protein–term networks of mPFC–BLA-enriched (**a**) and BLA–mPFC-enriched (**b**) proteins were constructed using KEGG BRITE annotations. Nodes represent proteins and functional terms, with edges indicating associations based on BRITE functional classifications. Note that mPFC–BLA-enriched proteins in this figure are defined as those uniquely identified in the mPFC–BLA neural pathway, with no overlap with proteins enriched in the BLA–mPFC neural pathway.

**Supplemental Figure 2.**
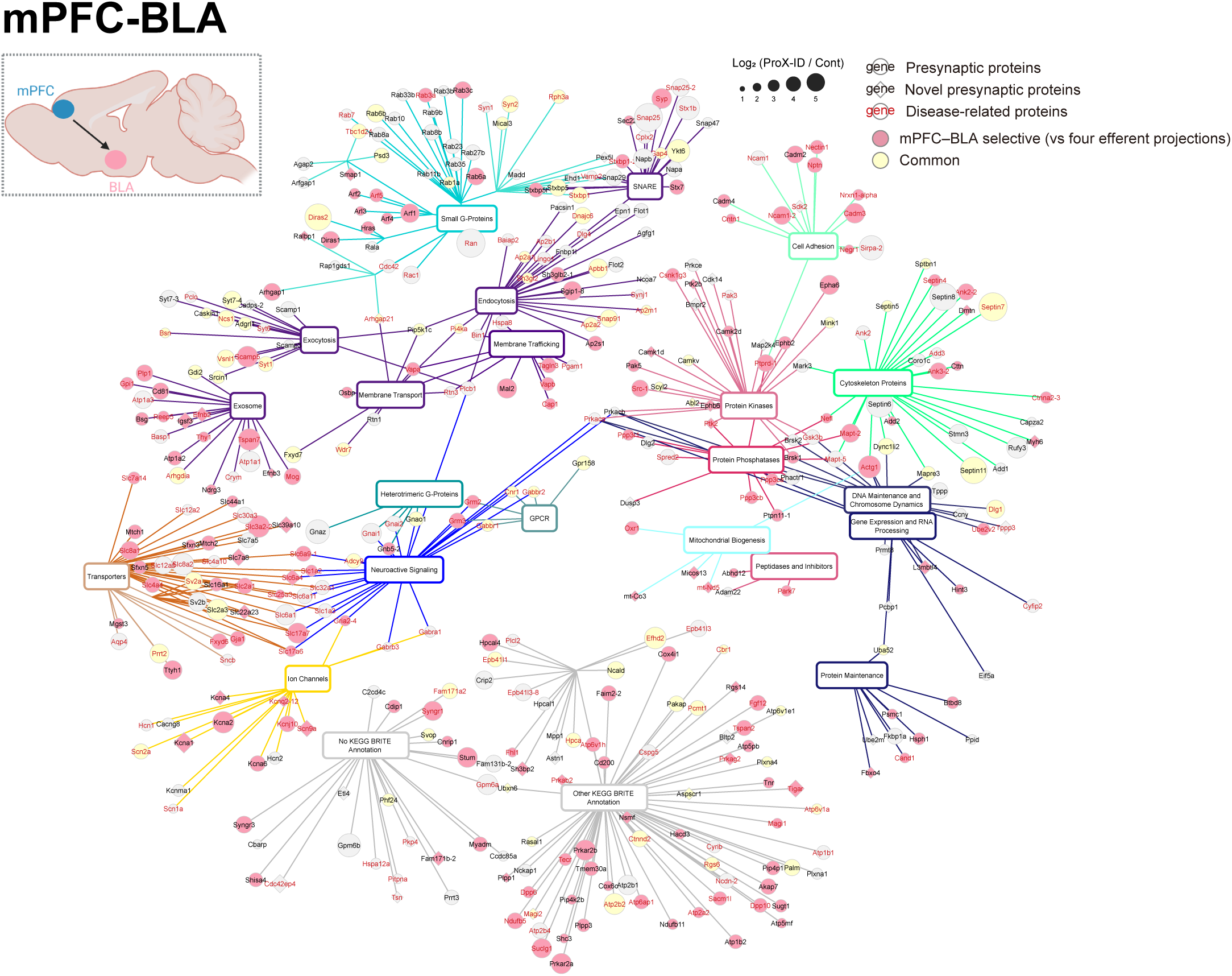
Protein–term network for mPFC–BLA-enriched neural pathway-enriched proteins. Protein–term networks were constructed using KEGG BRITE annotations. Nodes represent proteins and functional terms, with edges indicating associations based on BRITE functional classifications. Note that the mPFC–BLA-enriched proteins in this figure are defined as those uniquely identified in the mPFC–BLA neural pathway, with no overlap with proteins from the other four mPFC neural pathways.

**Supplemental Figure 3.**
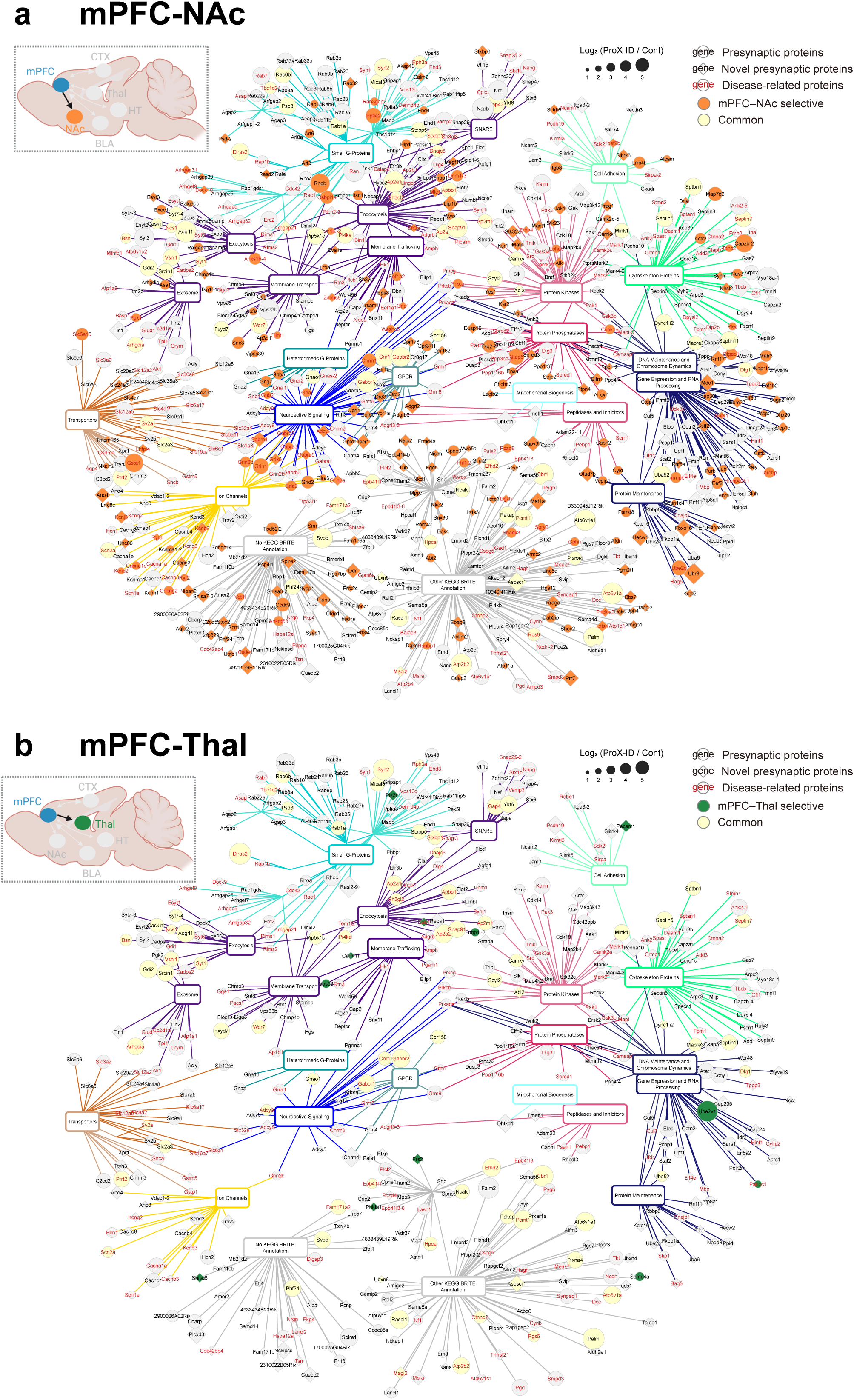
Protein–term network for neural pathway-enriched proteins. Protein–term networks for mPFC–NAc-enriched (**a**) and mPFC–Thal–enriched (**b**) proteins were constructed using KEGG BRITE annotations. Nodes represent proteins and functional terms, with edges indicating associations based on BRITE functional classifications.

**Supplemental Figure 4.**
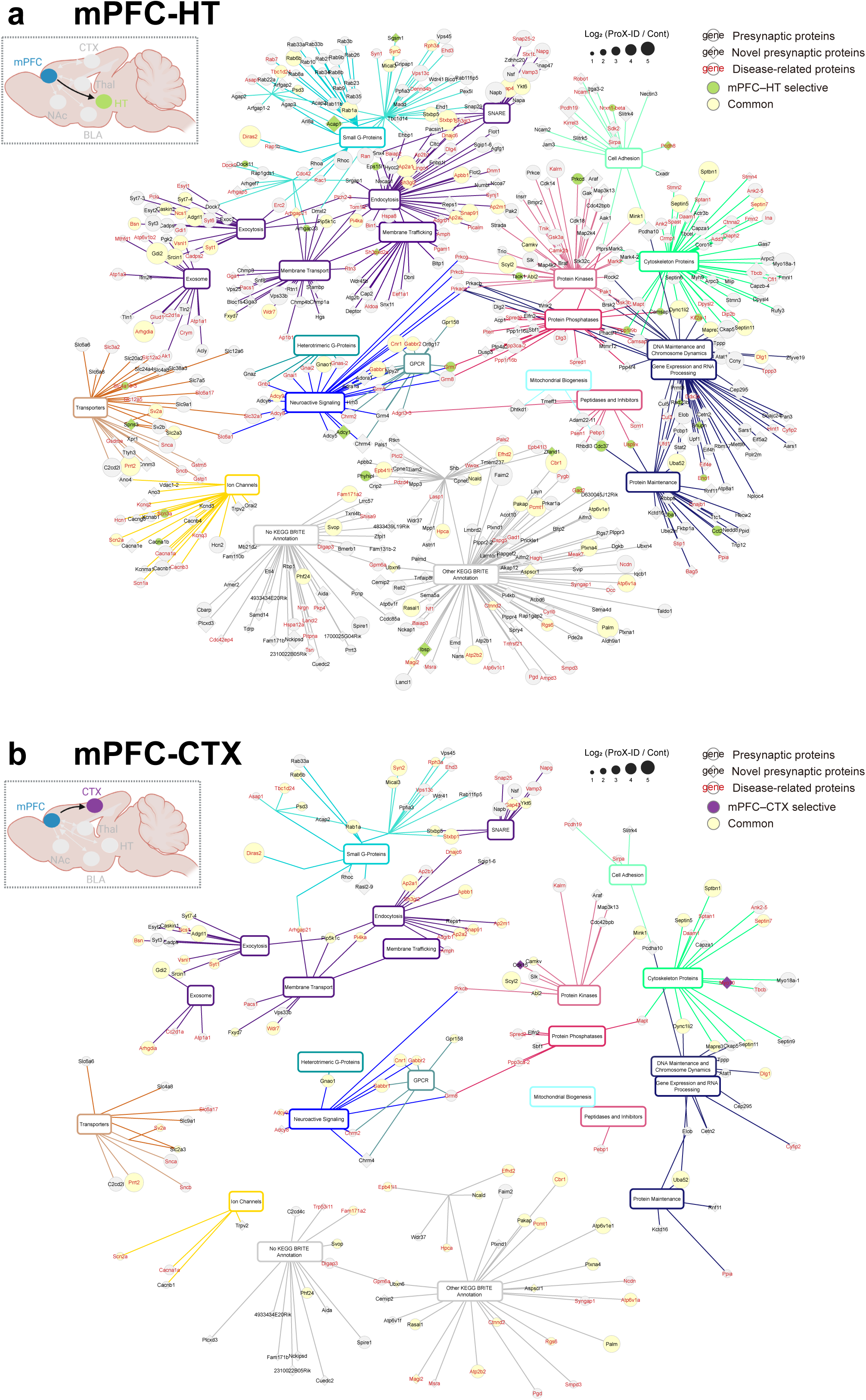
Protein–term network for neural pathway-enriched proteins. Protein–term networks for mPFC–HT-enriched (**a**) and mPFC–CTX-enriched (**b**) proteins were constructed using KEGG BRITE annotations. Nodes represent proteins and functional terms, with edges indicating associations based on BRITE functional classifications.

**Supplemental Figure 5.**
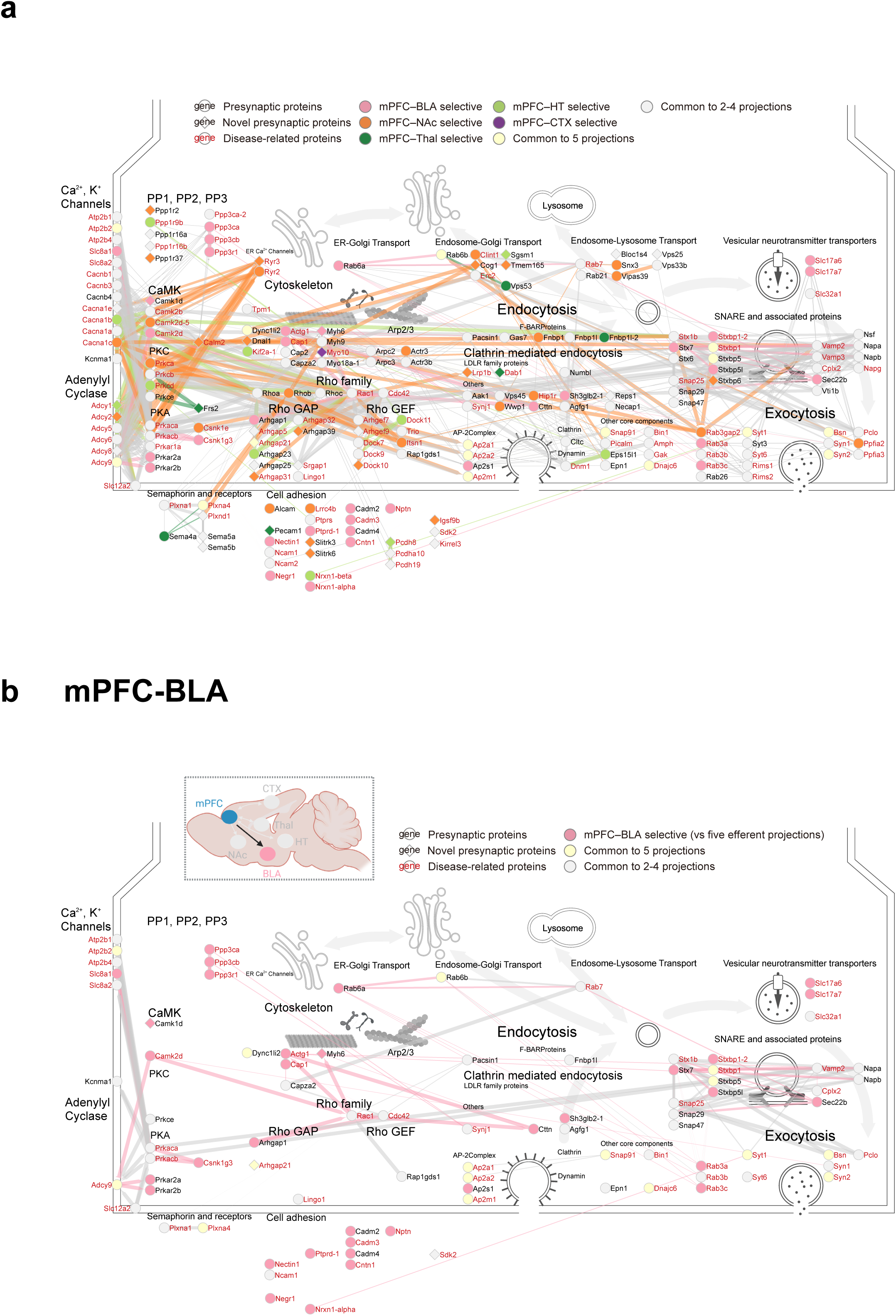
Neural pathway-enriched proteins involved in representative presynaptic events. Proteins enriched in at least one of the five mPFC efferent pathways (**a**) and mPFC–BLA-enriched proteins (**b**) involved in representative presynaptic events. Edges represent protein–protein interactions (PPIs) retrieved using the Reactome FI plugin in Cytoscape. Edge thickness and transparency reflect FI scores, which range from 0.89 to 1. For visualization, the edge thickness and transparency of PPIs annotated as “complex” were displayed using the same line style as those with an FI score of 0.1.

**Supplemental Figure 6.**
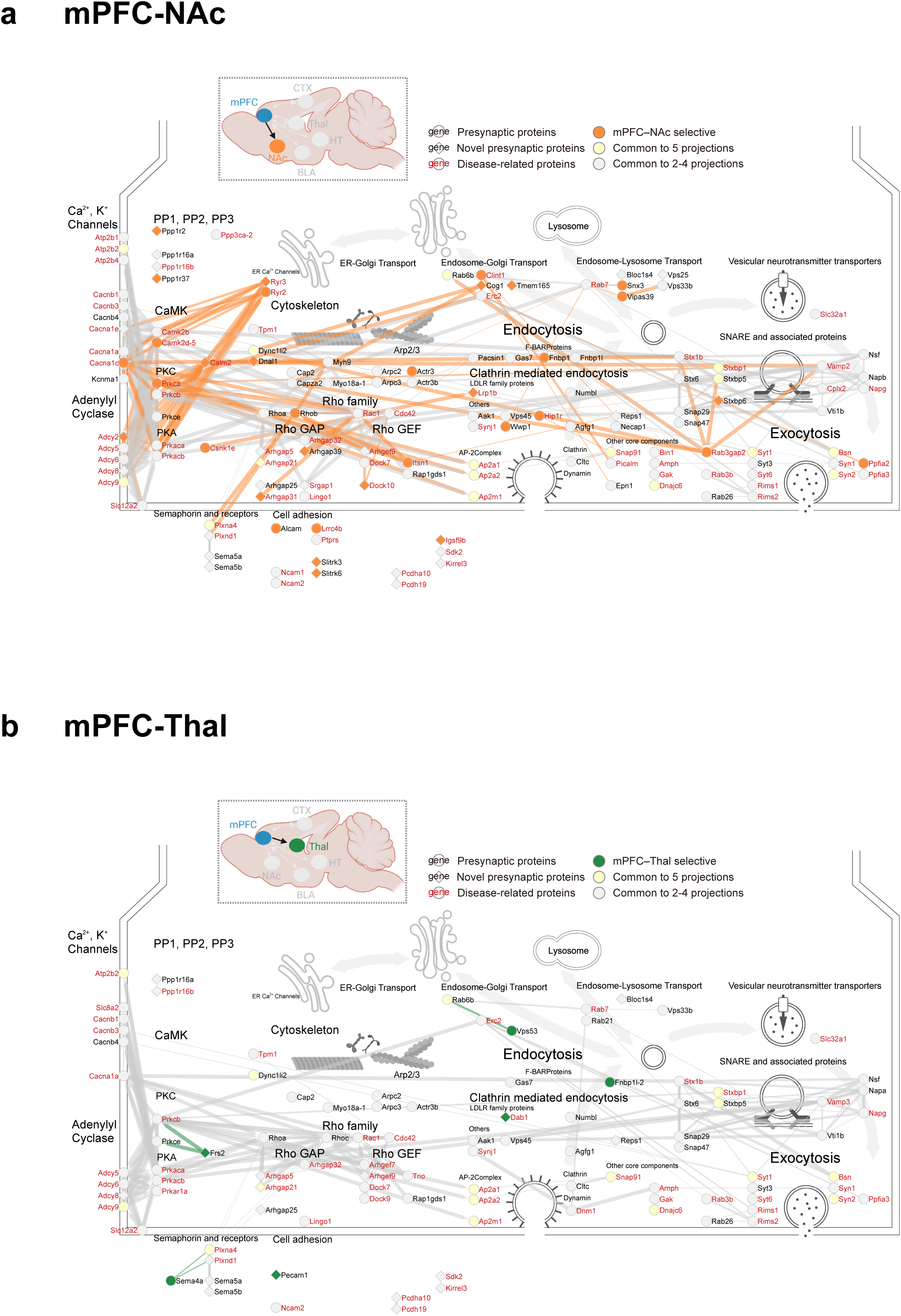
Neural pathway-enriched proteins involved in representative presynaptic events. mPFC–NAc-enriched (**a**) and mPFC–Thal-enriched (**b**) proteins involved in representative presynaptic events. Edges represent protein–protein interactions (PPIs) retrieved using the Reactome FI plugin in Cytoscape. Edge thickness and transparency indicate FI scores, which range from 0.89 to 1. For visualization, the edge thickness and transparency of PPIs annotated as “complex” were displayed using the same line style as those with an FI score of 0.1.

**Supplemental Figure 7.**
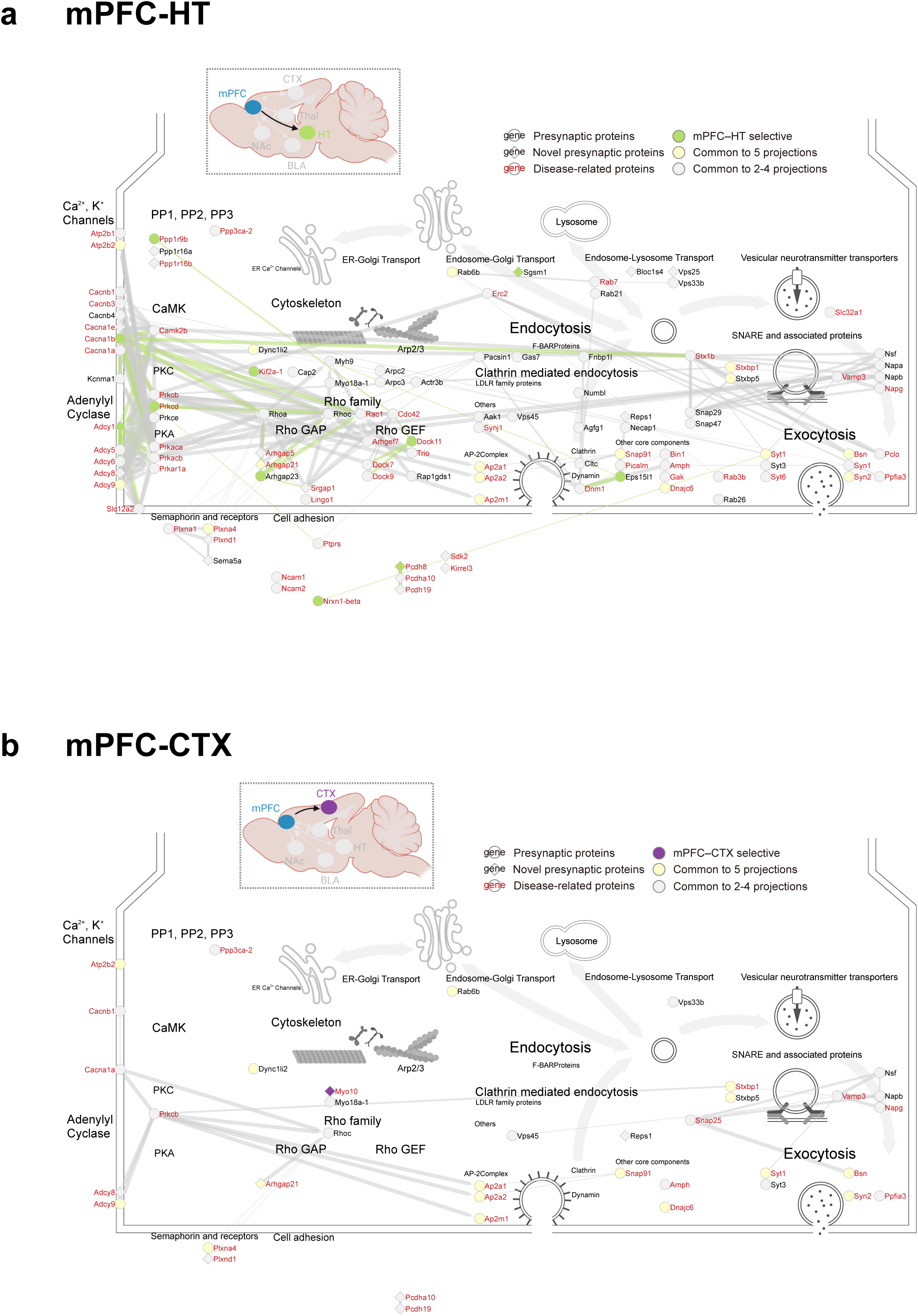
Neural pathway-enriched proteins involved in representative presynaptic events. mPFC–HT-enriched (**a**) and mPFC–CTX-enriched (**b**) proteins involved in representative presynaptic events. Edges represent protein–protein interactions (PPIs) retrieved using the Reactome FI plugin in Cytoscape. Edge thickness and transparency indicate FI scores, which range from 0.89 to 1. For visualization, the edge thickness and transparency of PPIs annotated as “complex” were displayed using the same line style as those with an FI score of 0.1.

**Supplemental Figure 8.**
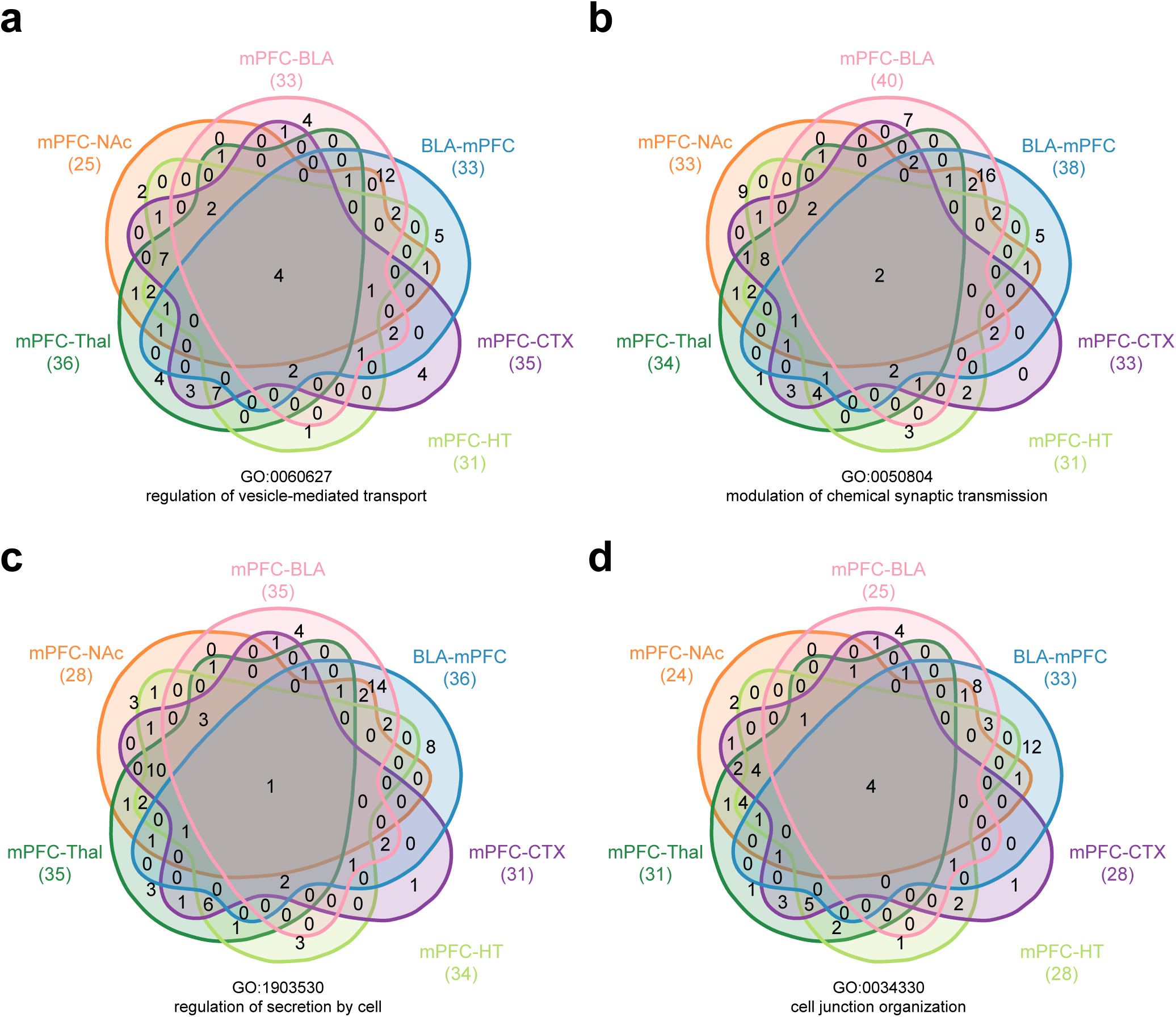
**a**–**d,** Venn diagrams showing the overlap of neural pathway-enriched proteins among the six neural pathways for representative GO terms. Numbers indicate the number of proteins in each intersection. The numbers in parentheses below each projection indicate the number of proteins annotated in the GO terms.

**Supplemental Figure 9.**
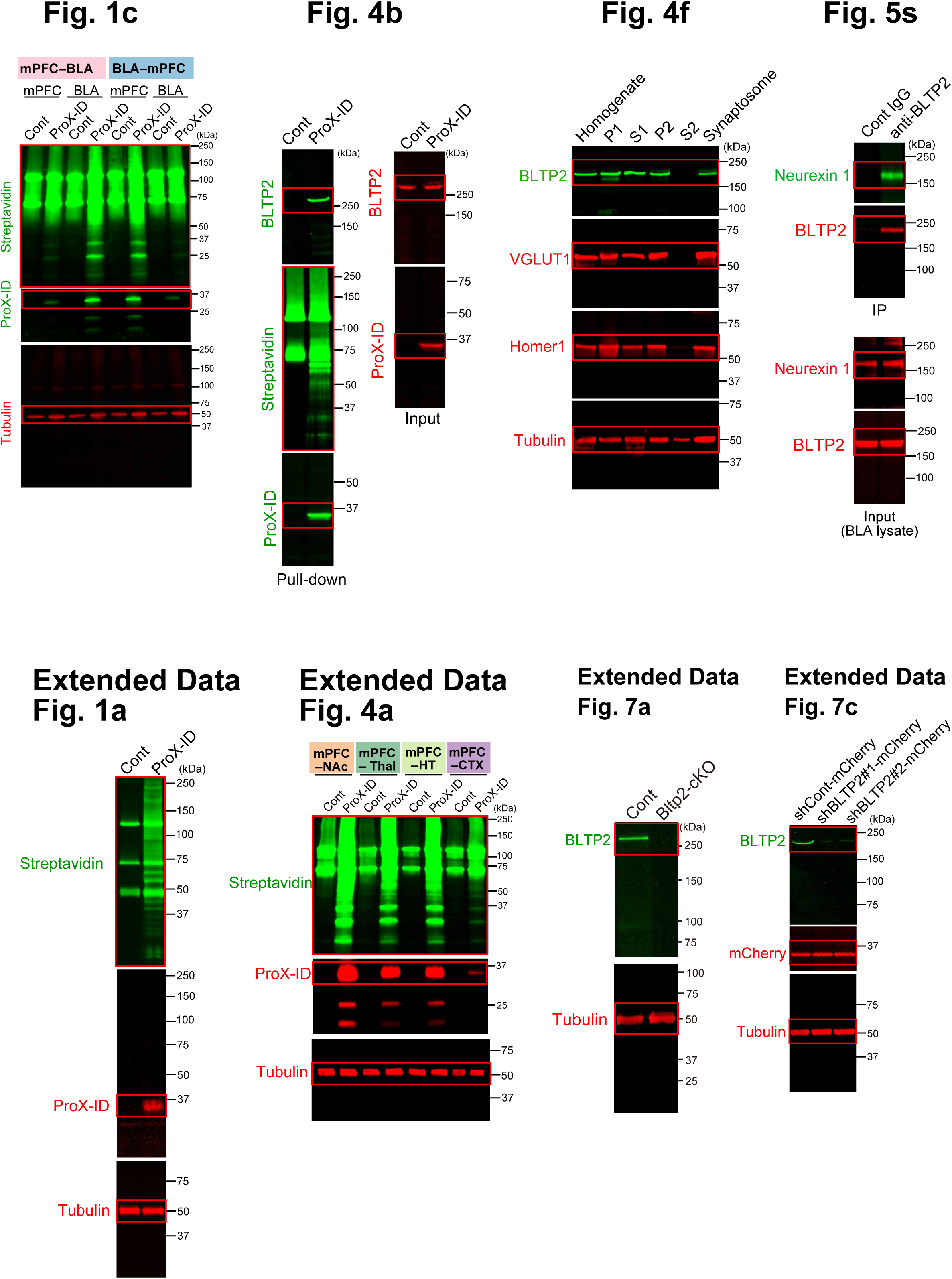
Full-size images of immunoblots.

